# Memory T Cells in MHC-Deficient Humanized Mice

**DOI:** 10.64898/2026.08.20.745695

**Authors:** M. Dargužytė, Z. Zhumadilova, F. Khan, M. Rahman, Sagar, A. Ernst, I. Poschke, J. Schulte-Schrepping, E. De-Domenico, MD Beyer, D. Schaudien, A. Dragon, B. Eiz-Vesper, C. von Kaisenberg, F. Klawonn, M. Thelen, H. Schlößer, E. Bauer, F. Klein, A. Schmitt, B. Soper, LD Shultz, R. Stripecke

**Affiliations:** Institute for Translational Immune-Oncology, Cancer Research Center Cologne-Essen (CCCE), Clinic of Internal Medicine I, University Hospital Cologne, University of Cologne, Cologne, Germany; Center for Molecular Medicine Cologne (CMMC) and Translational Center for Infectious Diseases and Oncology (TRIO), University of Cologne, Faculty of Medicine and University Hospital Cologne, Cologne, Germany; Department of Medicine II (Gastroenterology, Hepatology, Endocrinology, and Infectious Diseases), Freiburg University Medical Center, Faculty of Medicine, University of Freiburg, Freiburg, Germany; Faculty of Biology, University of Freiburg, Freiburg, Germany; Immune Monitoring Unit, National Center for Tumor Diseases Heidelberg, Heidelberg, Germany; Clinical Cooperation Unit Neuroimmunology and Brain Tumor Immunology, German Cancer Research Center, Heidelberg, Germany; German Cancer Consortium (DKTK), DKFZ core center, Heidelberg, Germany; Platform for Single Cell Genomics and Epigenomics (PRECISE), Deutsches Zentrum für Neurodegenerative Erkrankungen (DZNE) and University of Bonn and West German Genome Center, Bonn, Germany; Genomics and Immunoregulation, Life & Medical Sciences (LIMES) Institute, University of Bonn, Bonn, Germany; Immunogenomics & Neurodegeneration, Deutsches Zentrum für Neurodegenerative Erkrankungen (DZNE), Bonn, Germany; Fraunhofer Institute for Toxicology and Experimental Medicine ITEM, Hannover, Germany; Institute of Transfusion Medicine and Transplant Engineering, Hannover Medical School, Hannover, Germany; Clinic of Gynecology and Obstetrics, Hannover Medical School, Hannover, Germany; Department of Computer Science, Ostfalia University of Applied Sciences, Wolfenbuettel, Germany; Department of General, Visceral, Thoracic, and Transplantation Surgery, Faculty of Medicine and University Hospital Cologne, University of Cologne, Cologne, Germany; Institute of Transfusion Medicine, Faculty of Medicine and University Hospital Cologne, University of Cologne, Cologne, Germany; Laboratory of Experimental Immunology, Institute of Virology, Faculty of Medicine and University Hospital Cologne, University of Cologne, Cologne, Germany; The Jackson Laboratory, Bar Harbor, USA; Clinic of Hematology, Oncology, Hemostasis and Cell Therapy, Hannover Medical School, Hannover, Germany

**Keywords:** Humanized mice, T cell, MHC, single-cell mRNA, preclinical models

## Abstract

Major histocompatibility complexes (MHC) govern antigen presentation and T-cell receptor (TCR) selection. Accurate *in vivo* modeling of human immunity therefore requires physiological human MHC–TCR interactions. Humanized NOD-scid-IL2Rγc^null^ (NSG) mice engrafted with human CD34⁺ hematopoietic stem cells are widely used to provide preclinical platforms for the development of advanced therapies; however, interactions between murine MHC and human TCR can promote xenoreactivity and alter T-cell development. Here, we investigated how elimination of murine MHC together with different conditioning regimens shapes human T-cell maturation *in vivo*. CD34⁺ cells from ten cord blood donors were transplanted into conventional NSG mice or murine MHC-deficient NSG derivatives (DKO) following either sublethal irradiation or myeloablative busulfan conditioning. Integrated analyses combining flow cytometry, plasma cytokine profiling, and bulk and single-cell TCR sequencing revealed marked differences in T-cell differentiation across models. Busulfan-conditioned DKO mice developed highly proliferative, activated, and cytotoxic T cells together with clonally expanded TCR repertoires. In contrast, irradiated NSG mice preferentially accumulated naïve, NKT, and regulatory T-cell populations. Busulfan-conditioned DKO mice showed no evidence of xenogeneic graft-versus-host disease and represent a refined enabling platform for human T-cell development and provide a foundation for future preclinical evaluation of advanced gene and cell therapies.

## Introduction

Mice with a human immune system (HIS) - immunodeficient murine recipients engrafted with human hematopoietic stem and progenitor cells (HSPC) and/or lymphoid tissues - have become critical tools for modeling human immunity *in vivo*. These models enable mechanistic studies of human hematopoiesis, lymphocyte development, infections, immunopathologies, and human responses to vaccines and immunotherapies (1–5). Generation of the severe combined immune deficiency (SCID) and non-obese diabetic (NOD) strain carrying the IL2rγ-null mutation (NOD-SCID-gamma/NSG) enabled high levels of human lymphoid cell engraftment, permitting multilineage long-term reconstitution of human T cells, B cells, and innate subsets such as monocytes and NK cells in peripheral tissues (6,7). A central overall challenge in the conventional HIS preclinical models using different immunodeficient mouse strains is the occasional xenoreactivity caused by murine major histocompatibility complex (MHC) expression (8).

A major limitation of conventional humanized immune system (HIS) mouse models is that human T cells develop within a suboptimal immune microenvironment. In immunocompetent mice, the thymus is the principal site of T-cell development, where positive and negative selections are mediated largely by murine MHC molecules expressed on thymic epithelial and dendritic cells. However, extrathymic T-cell development may also contribute under conditions of thymic hypoplasia, lymphopenia, or following hematopoietic transplantation of humanized NSG mice. Consequently, the human TCR repertoire is shaped predominantly in the context of murine rather than human MHC (human leukocyte antigen, HLA) molecules. This may bias T-cell responses toward murine MHC–peptide complexes while limiting optimal recognition of antigens presented by autologous human HLA-expressing antigen-presenting, infected, or tumor cells. As these T cells retain functional endogenous TCRs, recognition of murine MHC expressed in host tissues can promote xenogeneic T-cell activation, inflammatory cytokine production, weight loss, and graft-versus-host disease.

To overcome this limitation, genetic ablation of murine MHC class I and II molecules has been combined with transgenic expression of defined human HLA alleles. When human HSPC donors are HLA-matched to these recipient strains, HLA-restricted T-cell development, antigen-specific antiviral immunity, and B-cell class-switch recombination are significantly improved compared with conventional humanized mouse models (9–11).

Further advancement of less complex HIS models that bypass the need for graft selection based on polymorphic HLA matching - while minimizing interference between murine MHC and human HLA molecules and maintaining responsiveness across diverse HLA backgrounds - is critical for accurate preclinical investigation of antigen-specific T cell responses to infectious diseases and malignancies, as well as for the evaluation of immunotherapies and studies of human immune biology.

We previously evaluated long-term (20-week) human hematopoiesis and T cell development in NSG mice engineered with the MHC class I and II double knockout (DKO) (12). Using CD34^+^ isolated HSPC from three cord-blood donors, stable human immune reconstitution in peripheral blood was seen from 8 to 20 weeks post-transplant, with single-positive CD4⁺ and CD8⁺ T cells detectable in thymus, bone marrow, and spleen. Systemic delivery of a lentiviral vector (LV) encoding HLA-A*02:01, HLA-DRB1*04:01, human GM-CSF/IFN-α, and cytomegalovirus gB antigen enhanced the presence of αβ and γδ T cells, NK cells, and HLA-DR-expressing lymphocytes in bone marrow. Transcriptomic analysis revealed that LV delivery upregulated pathways associated with antiviral and broader defense responses, indicating that DKO mice support functional, antigen-responsive human lymphocytes capable of mounting adaptive and innate immune programs *in vivo*. A major limitation of our previous study characterizing the mouse MHC interference in human T cell development in the DKO model was the lack of a side-by-side comparison with the parental NSG strain. In addition, we observed that the DKO strain was highly susceptible to irradiation, resulting in delayed weight gain.

Here, we therefore conducted a comprehensive comparison of NSG and DKO mice transplanted side-by-side with HSPC from ten individual cord-blood donors. To reduce irradiation-associated toxicity in DKO mice, we evaluated myeloablative conditioning with busulfan prior to transplantation as an alternative preconditioning strategy. DKO mice exhibited enhanced development of human memory T cells, characterized by a more clonally restricted TCR repertoire, whereas NSG mice predominantly generated naïve T cells with lower clonality. Moreover, busulfan preconditioning markedly improved the expansion and differentiation of human T cells with cytotoxic features. These findings indicate that murine MHC deficiency, particularly when combined with optimized conditioning, promotes more mature and functionally specialized human T cell responses in humanized mice.

## Results

### Busulfan preconditioning of DKO mice is associated with improved body weight recovery and supports robust CD4/CD8 T cell development

In a previous study, we demonstrated that DKO mice could be successfully humanized with isolated cord blood (CB) - mononuclear (MC) CD34^+^ cells following irradiation preconditioning (12). To directly assess the impact of the mouse host strain and preconditioning regimen on humanization efficiency, we performed a head-to-head comparison of DKO mice and their parental NSG strain. 6-week-old female mice were preconditioned with either X-ray irradiation (IR) or myeloablative busulfan (BU) treatment prior to hematopoietic cell transplantation (HCT) (Figure 1A). We utilized CD34^+^ cells from ten CB donors representing diverse HLA haplotypes (Table S1). Each CB unit was distributed across four experimental cohorts: DKO-BU, DKO-IR, NSG-BU, and NSG-IR. Humanized mice were monitored for 20 weeks post-HCT, with body weights recorded every two days and mice monitored for signs of graft-versus-host disease (GvHD) or adverse effects. Peripheral blood (PBL) samples were collected on weeks 8, 12, and 16 after transplantation. Mice were humanely euthanized on week 20 for tissue collection. Busulfan-treated mice demonstrated superior weight gain compared to irradiated mice across both strains (Figure 1B, for each mouse used in cohorts, see Figure S1A; mean weight increase from day -1 to day 138 post-HCT: DKO-BU 6.1 ± 1.5 g, DKO-IR 4.6 ± 1.5 g, NSG-BU 6.6 ± 0.7 g, NSG-IR 5.8 ± 1.5 g). A subset of DKO-IR mice (4/10) developed transient focal alopecia on the dorsal head region, observable between 30-90 days post-irradiation. One mouse each from the DKO-IR and NSG-IR cohorts died of undetermined causes during the observation period. Post-mortem analyses were not possible because the deaths occurred sporadically during the study, and the carcasses were not preserved for subsequent pathological examination. Therefore, overall, busulfan conditioning was associated with improved body weight recovery after HCT compared with irradiation under our experimental conditions.

**Figure 1.**
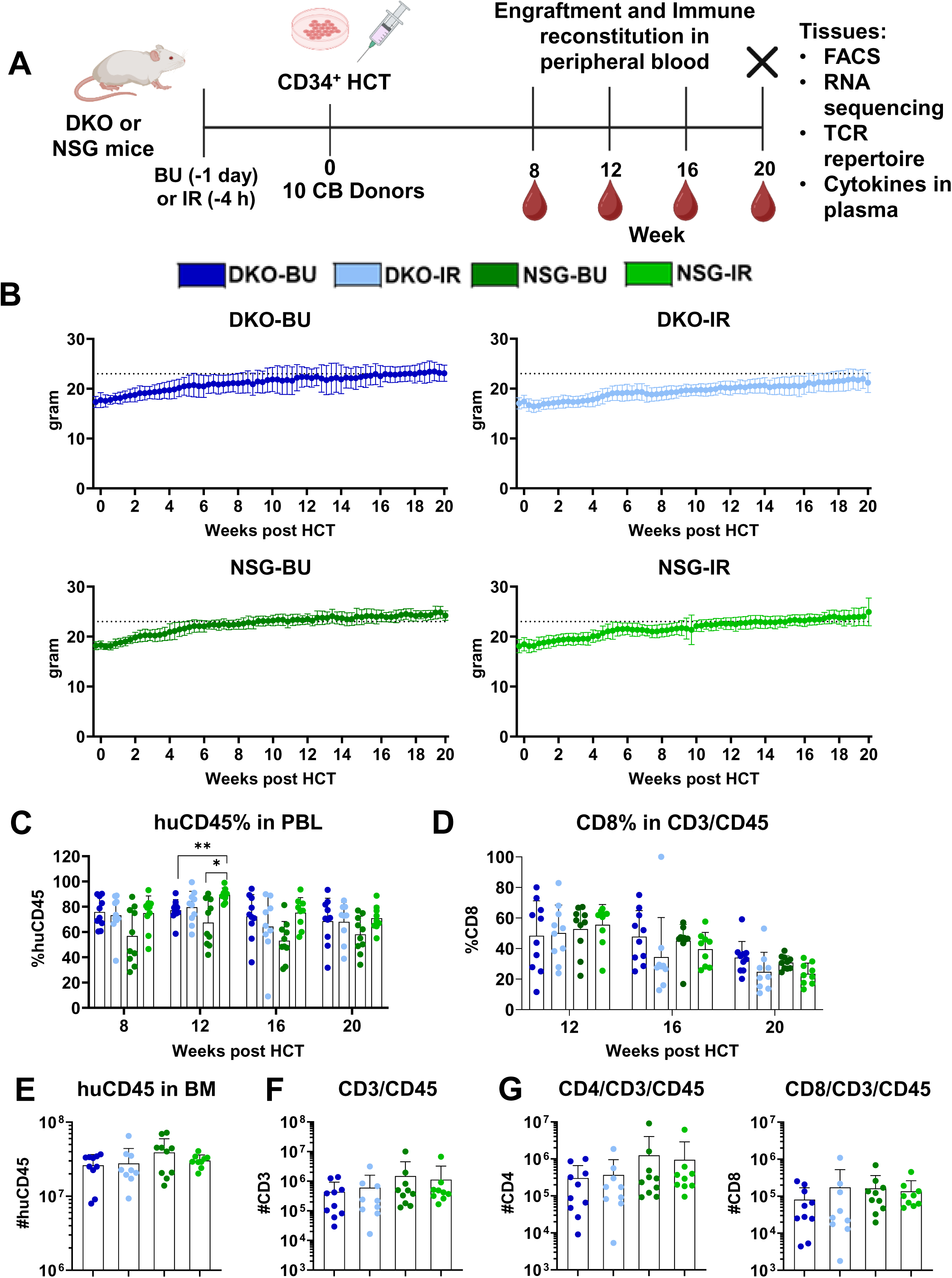
Comparative analyses of weight, long-term human hematopoietic engraftment, and immune reconstitution. A. Experimental scheme. Mice were either irradiated or received busulfan as preconditioning before CD34^+^ stem cell transplantation (HCT). The human immune reconstitution and engraftment was monitored after blood draws at weeks 8, 12, 16, and 20 post-HCT. After euthanasia, tissues were collected for FACS, RNA sequencing, TCR repertoire analyses, and detection of human cytokines in plasma. Humanized DKO mice preconditioned with busulfan or with irradiation were compared in parallel with humanized NSG mice preconditioned with busulfan or with irradiation. One DKO-IR and one NSG-IR mouse succumbed during the experiment. B. Average weight change curves over the course of the experiment in grams. Humanized DKO-BU mice (dark blue) show higher weight gain compared to DKO-IR (light blue). A similar trend is seen in NSG mice between two preconditioning protocols. C. Analyses of huCD45^+^ lymphocytes frequencies in peripheral blood (PBL) at weeks 8, 12, 16, and 20 after HCT (in percentages). D. Analyses of CD8^+^ cell frequencies within CD3/CD45 in blood at weeks 12, 16, and 20 after HCT (in percentages). E. Quantified total huCD45 cell counts in bone marrows at week 20 post HCT (in log scale). F. Total CD3^+^ cell counts in bone marrow (in log scale). G. Total CD4^+^ or CD8^+^ T cell counts in bone marrow (in log scale). DKO-BU (dark blue), DKO-IR (light blue), NSG-BU (dark green), and NSG-IR (light green). Data are presented as mean and SD, and individual values are shown. The differences were analyzed with the Wilcoxon test with Bonferroni correction (n = 9 - 10). **** = p < 0.0001; *** = p < 0.001; ** = p < 0.01; * = p < 0.05.

Despite variations in HLA genotype, mouse strain, and preconditioning regimen, human CD45^+^ (huCD45^+^) cell frequencies in peripheral blood ranged between 60-90% from weeks 8-20 post-HCT across most cohorts, with the NSG-BU group showing lower average engraftment than NSG-IR, which was significant on week 12 post HCT (Table 1, Figure 1C). Human CD3^+^ T cell frequencies in blood increased progressively, reaching approximately 30% by week 20 post-HCT (Figure S1B; DKO-BU: 28.3 ± 19.7%, DKO-IR: 24.6 ± 19.2%, NSG-BU: 27.8 ± 21.7%, NSG-IR: 34.8 ± 21.3%). Within the T cell compartment, CD8^+^ T cell frequencies declined over time across all cohorts but remained highest in DKO-BU mice at week 20 post-HCT (Figure 1D). Conversely, CD4^+^ T cell frequencies increased during the course of the experiment and were lowest in DKO-BU mice and significantly elevated in the NSG-IR cohort (Figure S1C). Long-term huCD45^+^ hematopoietic engraftment in bone marrow was robust across all cohorts (Figure 1E; DKO-BU: 2.6 × 10^7^ ± 1.0 × 10^7^, DKO-IR: 2.8 × 10^7^ ± 1.6 × 10^7^, NSG-BU: 3.9 × 10^7^ ± 2.1 × 10^7^, NSG-IR: 3.0 × 10^7^ ± 0.6 × 10^7^). Total CD3^+^ T cell numbers within the huCD45^+^ population in bone marrow were lower in DKO mice compared to NSG mice across both preconditioning regimens (Figure 1F; DKO-BU: 0.5 × 10^6^ ± 0.5 × 10^6^, DKO-IR: 0.6 × 10^6^ ± 1.0 × 10^6^, NSG-BU: 1.5 × 10^6^ ± 3.0 × 10^6^, NSG-IR: 1.1 × 10^6^ ± 2.1 × 10^6^). CD4^+^CD3^+^ T cells predominated over CD8^+^CD3^+^ T cells in bone marrow across all experimental groups (Figure 1G; for percentages see Figure S1D). Collectively, these findings demonstrate that DKO mice tolerate busulfan preconditioning better than irradiation while maintaining robust humanization and T cell development overall comparable to NSG controls.

**Table 1:** Detection of human hematopoietic cells in peripheral blood over time. Flow cytometry analyses were performed at weeks 8, 12, 16, and 20 after stem cell transplantation of DKO (A) or NSG (B) mice receiving busulfan (BU) or irradiation (IR) preconditioning.

|  | huCD45% at week 8 |  | huCD45% at week 12 |  | huCD45% at week 16 |  | huCD45% at week 20 |  |
| --- | --- | --- | --- | --- | --- | --- | --- | --- |
| Cord Blood ID | DKO BU | DKO IR | DKO BU | DKO IR | DKO BU | DKO IR | DKO BU | DKO IR |
| #74 | 78.71 | 83.99 | 79.44 | 82.59 | 54.78 | 52.56 | 81.75 | 61.35 |
| #96 | 71.27 | 70.58 | 79.26 | 71.11 | 73.87 | n.a. | 69.75 | n.a. |
| #119 | 90.30 | 88.42 | 76.00 | 85.94 | 94.23 | 78.72 | 93.36 | 79.98 |
| #121 | 67.07 | 66.47 | 73.13 | 74.07 | 71.81 | 64.93 | 83.97 | 72.88 |
| **#160** | 89.94 | 88.87 | 90.78 | 91.06 | 86.90 | 75.64 | 77.32 | 73.47 |
| **#161** | 68.85 | 82.85 | 83.68 | 87.66 | 89.24 | 58.16 | 49.12 | 73.11 |
| **#189** | 88.40 | 73.59 | 74.74 | 64.92 | 70.93 | 61.21 | 56.27 | 48.53 |
| **#190** | 60.74 | 37.25 | 75.44 | 81.42 | 79.29 | 88.86 | 70.68 | 79.71 |
| **#312** | 82.89 | 69.40 | 81.55 | 99.49 | 66.79 | 89.93 | 71.13 | 84.96 |
| **#342** | 60.46 | 70.78 | 58.61 | 58.05 | 36.01 | 9.04 | 31.78 | 38.81 |
| **Average**<br>**± SD** | 75.86 ±<br>11.71 | 73.22 ±<br>15.10 | 77.26 ±<br>8.35 | 79.63 ±<br>12.59 | 72.39 ±<br>17.28 | 64.34 ±<br>24.59 | 68.51 ±<br>18.25 | 68.09 ±<br>15.51 |
| **Cord**<br>**Blood ID** | **NSG BU** | **NSG IR** | **NSG BU** | **NSG IR** | **NSG BU** | **NSG IR** | **NSG BU** | **NSG IR** |
| **#74** | 59.86 | 82.41 | 58.53 | 87.18 | 47.55 | 76.92 | 62.14 | 64.54 |
| **#96** | 75.06 | 79.91 | 41.98 | 88.48 | 54.59 | 84.28 | 50.26 | 65.00 |
| **#119** | 38.89 | 75.09 | 85.97 | 98.92 | 81.31 | 93.65 | 77.03 | 79.31 |
| **#121** | 28.33 | 79.27 | 78.67 | 90.86 | 48.25 | 79.48 | 55.09 | 88.71 |
| **#160** | 87.89 | 80.51 | 75.24 | 91.17 | 59.64 | 82.60 | 71.11 | 76.85 |
| **#161** | 43.14 | 72.64 | 56.33 | 82.99 | 30.75 | 68.00 | 42.36 | 67.03 |
| **#189** | 32.67 | 93.19 | 90.35 | 90.29 | 48.50 | 56.17 | 48.83 | 55.08 |
| **#190** | 80.27 | 82.88 | 85.96 | 81.53 | 61.95 | 76.43 | 74.94 | 68.85 |
| **#312** | 77.39 | 56.93 | 50.64 | 93.96 | 65.94 | 60.07 | 65.37 | 72.86 |
| **#342** | 44.86 | 46.50 | 49.36 | n.a. | 32.52 | n.a. | 34.11 | n.a. |
| **Average**<br>**± SD** | 56.84 ±<br>21.93 | 74.93 ±<br>13.60 | 67.40 ±<br>17.94 | 89.49 ±<br>5.32 | 53.10 ±<br>15.21 | 75.29 ±<br>11.95 | 58.12 ±<br>14.38 | 70.91 ±<br>9.82 |

Table 1 A
|  | huCD45% at week 8 |  | huCD45% at week 12 |  | huCD45% at week 16 |  | huCD45% at week 20 |  |
| --- | --- | --- | --- | --- | --- | --- | --- | --- |
| Cord Blood ID | DKO BU | DKO IR | DKO BU | DKO IR | DKO BU | DKO IR | DKO BU | DKO IR |
| #74 | 78.71 | 83.99 | 79.44 | 82.59 | 54.78 | 52.56 | 81.75 | 61.35 |
| #96 | 71.27 | 70.58 | 79.26 | 71.11 | 73.87 | n.a. | 69.75 | n.a. |
| #119 | 90.30 | 88.42 | 76.00 | 85.94 | 94.23 | 78.72 | 93.36 | 79.98 |
| #121 | 67.07 | 66.47 | 73.13 | 74.07 | 71.81 | 64.93 | 83.97 | 72.88 |
| #160 | 89.94 | 88.87 | 90.78 | 91.06 | 86.90 | 75.64 | 77.32 | 73.47 |
| #161 | 68.85 | 82.85 | 83.68 | 87.66 | 89.24 | 58.16 | 49.12 | 73.11 |
| #189 | 88.40 | 73.59 | 74.74 | 64.92 | 70.93 | 61.21 | 56.27 | 48.53 |
| #190 | 60.74 | 37.25 | 75.44 | 81.42 | 79.29 | 88.86 | 70.68 | 79.71 |
| #312 | 82.89 | 69.40 | 81.55 | 99.49 | 66.79 | 89.93 | 71.13 | 84.96 |
| #342 | 60.46 | 70.78 | 58.61 | 58.05 | 36.01 | 9.04 | 31.78 | 38.81 |
| Average ± SD | 75.86 ± 11.71 | 73.22 ± 15.10 | 77.26 ± 8.35 | 79.63 ± 12.59 | 72.39 ± 17.28 | 64.34 ± 24.59 | 68.51 ± 18.25 | 68.09 ± 15.51 |

B
|  | huCD45% at week 8 |  | huCD45% at week 12 |  | huCD45% at week 16 |  | huCD45% at week 20 |  |
| --- | --- | --- | --- | --- | --- | --- | --- | --- |
| Cord Blood ID | NSG BU | NSG IR | NSG BU | NSG IR | NSG BU | NSG IR | NSG BU | NSG IR |
| #74 | 59.86 | 82.41 | 58.53 | 87.18 | 47.55 | 76.92 | 62.14 | 64.54 |
| #96 | 75.06 | 79.91 | 41.98 | 88.48 | 54.59 | 84.28 | 50.26 | 65.00 |
| #119 | 38.89 | 75.09 | 85.97 | 98.92 | 81.31 | 93.65 | 77.03 | 79.31 |
| #121 | 28.33 | 79.27 | 78.67 | 90.86 | 48.25 | 79.48 | 55.09 | 88.71 |
| #160 | 87.89 | 80.51 | 75.24 | 91.17 | 59.64 | 82.60 | 71.11 | 76.85 |
| #161 | 43.14 | 72.64 | 56.33 | 82.99 | 30.75 | 68.00 | 42.36 | 67.03 |
| #189 | 32.67 | 93.19 | 90.35 | 90.29 | 48.50 | 56.17 | 48.83 | 55.08 |
| #190 | 80.27 | 82.88 | 85.96 | 81.53 | 61.95 | 76.43 | 74.94 | 68.85 |
| #312 | 77.39 | 56.93 | 50.64 | 93.96 | 65.94 | 60.07 | 65.37 | 72.86 |
| #342 | 44.86 | 46.50 | 49.36 | n.a. | 32.52 | n.a. | 34.11 | n.a. |
| Average ± SD | 56.84 ± 21.93 | 74.93 ± 13.60 | 67.40 ± 17.94 | 89.49 ± 5.32 | 53.10 ± 15.21 | 75.29 ± 11.95 | 58.12 ± 14.38 | 70.91 ± 9.82 |

### Busulfan-treatment reveals changes in distributions of mouse F4/80 positive myeloid cells and human CD4^+^ T cells in the spleen

Flow cytometric analysis of splenocytes harvested at week 20 post-HCT revealed modestly elevated numbers of human CD45^+^, CD3^+,^ and both CD4^+^ and CD8^+^ T cells in BU compared to IR mice (Figure 2A-C). Histological examination of spleen sections provided additional insights into cell morphologies across experimental cohorts. Hematoxylin and eosin (H&E) staining of paraffin-embedded spleen sections revealed clusters of morphologically enlarged cells in DKO mice (Figure 2D). Staining with F4/80 showed more intense detection of mouse macrophages surrounding these clusters in DKO mice. Human CD4 immunohistochemistry revealed enlarged CD4⁺ cells that frequently formed clustered structures in DKO mice. While activated CD4⁺ T cells are known to undergo characteristic morphological changes, including increased cell size, enhanced cytoplasmic volume, and an altered nuclear-to-cytoplasmic ratio, human CD4 is also expressed by subsets of myeloid cells. Therefore, the identity and activation status of these enlarged CD4⁺ cells cannot be conclusively determined by CD4 immunohistochemistry alone. To further address this question, additional staining for the human myeloid marker CD68 was performed. However, CD68 staining was generally sparse in both NSG and DKO tissues and did not support a definitive identification of the enlarged CD4⁺ cells as human myeloid cells. Thus, these findings are best interpreted as demonstrating the presence of prominent clusters of enlarged human CD4⁺ cells, often surrounded by murine macrophages, while the precise cellular identity of these enlarged cells remains to be determined by future studies using higher-resolution phenotypic approaches.

**Figure 2.**
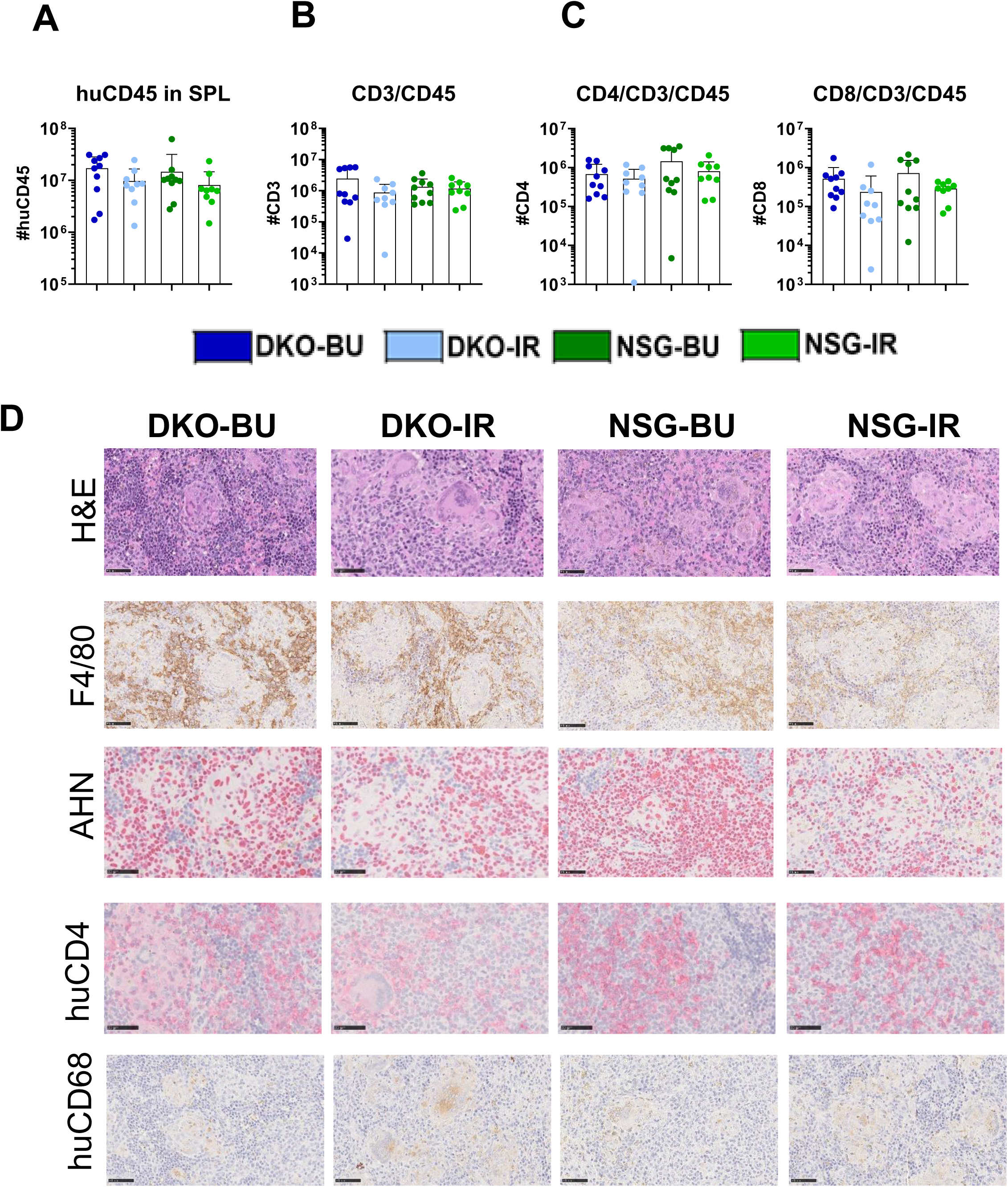
Analyses of spleens by flow cytometry and immunohistochemistry. A. Total huCD45 cell counts in splenocytes (in log scale). B. Total huCD3/CD45 cell counts in splenocytes (in log scale). C. Total huCD4/CD3/CD45 (left panel) and huCD4/CD3/CD45 (right panel) cell counts in splenocytes (in log scale). D. Representative images of spleens stained with H&E, or immunostained for detection of mouse macrophage marker (F4/80), human nucleus (AHN), human macrophage marker (CD68) and human CD4. The scale bars shown in the lower left corner of the pictures represent 50 µm. DKO-BU (dark blue), DKO-IR (light blue), NSG-BU (dark green), and NSG-IR (light green). Data are presented as mean and SD, and individual values are shown. The differences in A-C (n = 9 - 10) were analyzed with the Wilcoxon test with Bonferroni correction. **** = p < 0.0001; *** = p < 0.001; ** = p < 0.01; * = p < 0.05.

### Effector memory T cell frequencies are elevated in DKO mice, resembling adult human blood composition

Effector memory (EM) CD4^+^ and CD8^+^ T cells are characteristically more abundant in human peripheral blood mononuclear cells (PBMC) samples from adults compared to newborn CB-MC samples, reflecting progressive immune system maturation and antigen exposure over time (13). Conversely, CB-MC typically contains higher proportions of naïve (N) T cells, as the immune system remains in early developmental stages with ongoing thymic output. Comparisons of the T cell phenotypes by flow cytometry between cryopreserved CD34-negative fractions of CB-MC units used for mouse humanization and adult PBMC confirmed higher N frequencies in CB-MC and higher EM frequencies in PBMC (Figures 3A-D, Figure S2). Using the same methods, we then assessed the distribution of EM and N T cell subsets in freshly obtained spleen, peripheral blood, and bone marrow from humanized mice at week 20 post-HCT. The most pronounced differences emerged in splenic T cell populations. DKO mice exhibited more than twice as many and significantly higher frequencies of both CD4^+^ and CD8^+^ EM T cells compared to NSG mice (Figure 3A, C; CD8^+^ EM in spleen: DKO-BU 51.4 ± 8.2%, DKO-IR 46.6 ± 12.3%, NSG-BU 19.0 ± 12.5%, NSG-IR 23.6 ± 12.9%). Reciprocally, DKO cohorts contained significantly lower frequencies of CD4^+^ and CD8^+^ N T cells compared to NSG mice (Figure 3B, D). These contrasting memory versus naïve T cell distributions in DKO and NSG mice were consistently observed across PBL and bone marrow compartments. Central memory (CM) and terminally differentiated effector (TE) T cell frequencies showed lower variation among experimental cohorts (Figure S2). Collectively, these findings demonstrate that human T cells developing in DKO mice undergo enhanced maturation and activation compared to those in NSG mice, yielding T cell subset distributions that more closely resemble adult human PBMC rather than the naïve-predominant phenotype characteristic of CB-MC.

**Figure 3.**
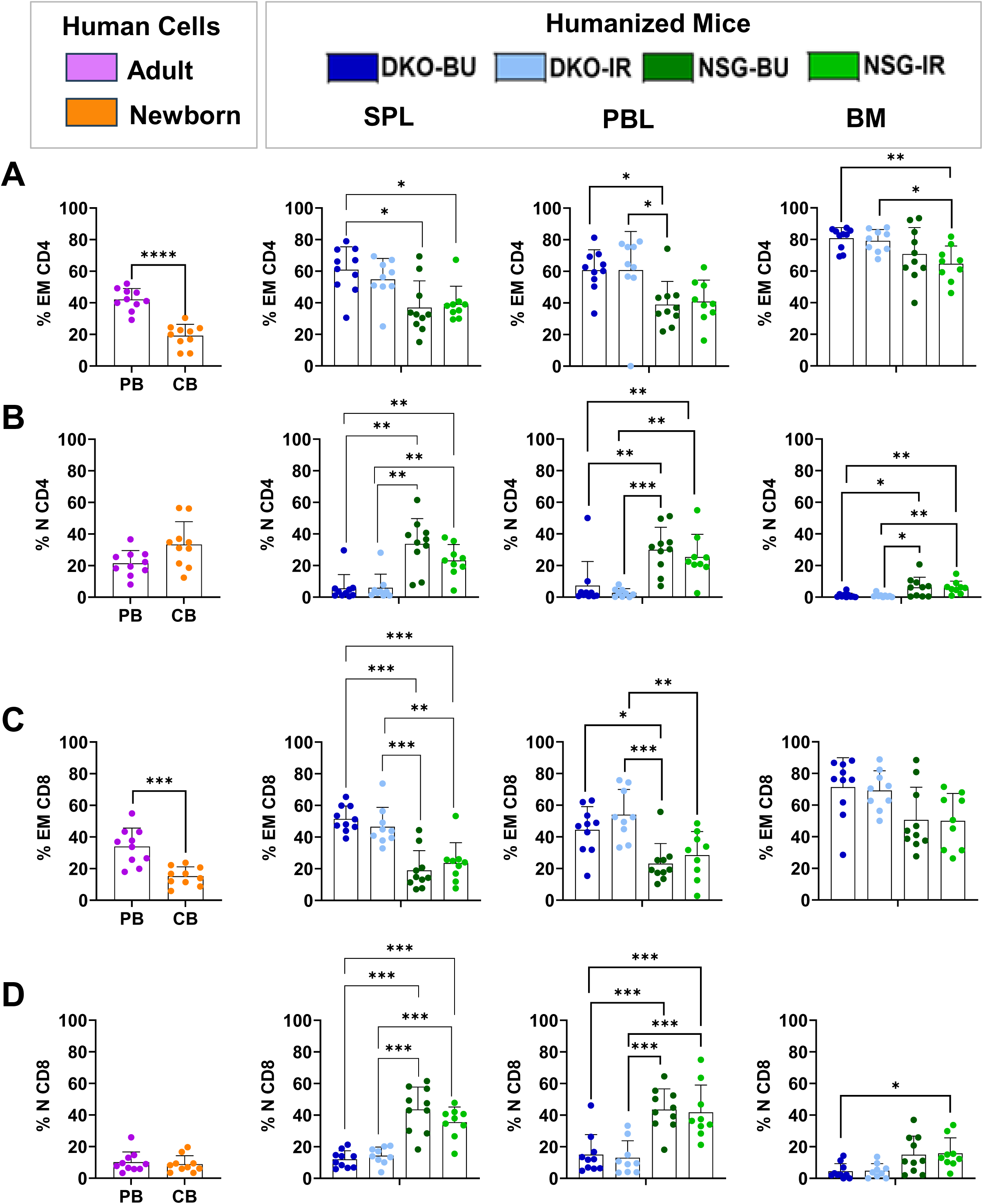
CD4+ and CD8+ T cell immunophenotype in adult PBMC (magenta), CD34-CB-MC (orange), and tissues of humanized mice (spleen /SPL, peripheral blood/ PBL, and bone marrow/ BM). DKO-BU (dark blue), DKO-IR (light blue), NSG-BU (dark green), and NSG-IR (light green). A. Frequencies of effector memory T cells within the CD4^+^ fraction (in percentage). B. Frequencies of naïve T cells within the CD4^+^ fraction (in percentage). C. Frequencies of effector memory T cells within the CD8^+^ fraction (in percentage). D. Frequencies of naïve T cells within the CD8^+^ fraction (in percentage). Data are presented as mean and SD, and individual values are shown. The differences were analyzed with the Wilcoxon test with Bonferroni correction (n = 9 – 10). **** = p < 0.0001; *** = p < 0.001; ** = p < 0.01; * = p < 0.05.

### Single-cell gene expression profiling identifies proliferative effector memory T cell clusters with cytotoxic properties in DKO-BU mice and naïve profiles in NSG mice

To comprehensively characterize T cell transcriptional states, we performed scRNA-seq analyses on splenic T cells from humanized mice (Figure 4A). We selected three representative CB donors that yielded distinct and unbiased T cell reconstitution patterns across mouse strains: CB74 (higher T cell counts in DKO mice), CB160 (comparable T cell counts between DKO and NSG mice), and CB161 (higher T cell counts in NSG mice) (Figure S3). CD3⁺ splenocytes were isolated from each cohort at week 20 post-HCT, and sequencing data were subsequently partitioned into CD4⁺ and CD8⁺ T cell subsets for independent analysis. Unsupervised clustering identified 13 distinct clusters of CD4⁺ T cells (Figure 4B). Clusters were manually annotated based on differential expression of canonical T cell subset markers (Figure 4C, S4A). Clustering of CD4⁺ T cells identified 13 clusters, which were grouped into seven functional subtypes. The Th1 subset (clusters 1 and 4) exhibited high expression of effector genes and pro-inflammatory cytokines, including CXCR3, TBX21, and IFNG (Figure S4A). Clusters 6, 10, and 11 represented proliferative cells marked by MKI67 (Figure S4A). The Th2-like subset (cluster 2) showed high GATA3 expression (Figure S4A). The Tfh subset (cluster 0) expressed BCL6, CXCR5, and IL21. Clusters 3, 5, 9, and 12 were classified as central memory (CM) CD4⁺ T cells, characterized by expression of naïve and migratory markers (TCF7, IL7R, SELL, and LEF1) (Figure S4A) and medium expansion TCR clones (Figure S5). The Treg subset (cluster 8) showed high expression of FOXP3 and CTLA4. Cluster 7 represented naïve CD4⁺ T cells with strong expression of migratory and naïve markers (CCR7, FOS, TCF7, IL7R, SELL, and LEF1). Clustering of unsupervised (Figure 4D) and annotated CD8⁺ T cells identified 12 clusters corresponding to nine subtypes (Figure 4E). Cluster 1, classified as cytotoxic T lymphocytes (CTLs), showed high expression of cytotoxic genes (GZMB, GNLY, and PRF1) (Figure S4B). Clusters 7, 8, and 9 were proliferative populations marked by MKI67. MAIT cells (cluster 5) exhibited high expression of lineage-associated genes, including KLRB1, TBX21, RORC, SLC4A10, and ITGAM (Figure S4B). Cluster 6, classified as CXCR6⁺ cells, expressed exhaustion and cytotoxic markers (CXCR6, LAG3, CTLA4, GZMB, and PRF1) (Figure S4B). Effector memory (EM) CD8⁺ T cells (cluster 0) showed high KLRG1 expression (Figure S4B) and were associated with medium-expansion TCR clones (Figure S5). Cluster 4, representing resident memory (RM) cells, displayed tissue residency signatures including KLRB1, ZNF683, and ITGAM. Naïve CD8⁺ T cells (clusters 2 and 3) expressed classical naïve and migratory markers (CCR7, FOS, TCF7, IL7R, SELL, and LEF1) (Figure S4B) and exhibited predominantly small expansion TCR clones with high repertoire diversity (Figure S5). Two additional minor populations were identified: a resident-like KLRD1⁺ subset (cluster 10) and NKT cells (cluster 11) (Figure S4B). We next examined the distribution of cells from each mouse cohort across the annotated clusters. Within the CD4⁺ T cell compartment, DKO mice predominantly harbored central memory and proliferative clusters, whereas NSG mice were enriched for naïve T cell clusters (Figure 4F, G). Similarly, within the CD8⁺ T cell compartment, DKO-derived T cells showed increased representation of proliferative and cytotoxic populations, while NSG mice displayed a predominance of naïve phenotype clusters (Figure 4H, I). Together, these single-cell transcriptomic analyses corroborate the flow cytometry findings and demonstrate at the transcriptional level that DKO mice support the development of activated, proliferative effector memory T cells with cytotoxic potential, whereas NSG mice predominantly maintain T cells in a transcriptionally naïve pattern.

**Figure 4.**
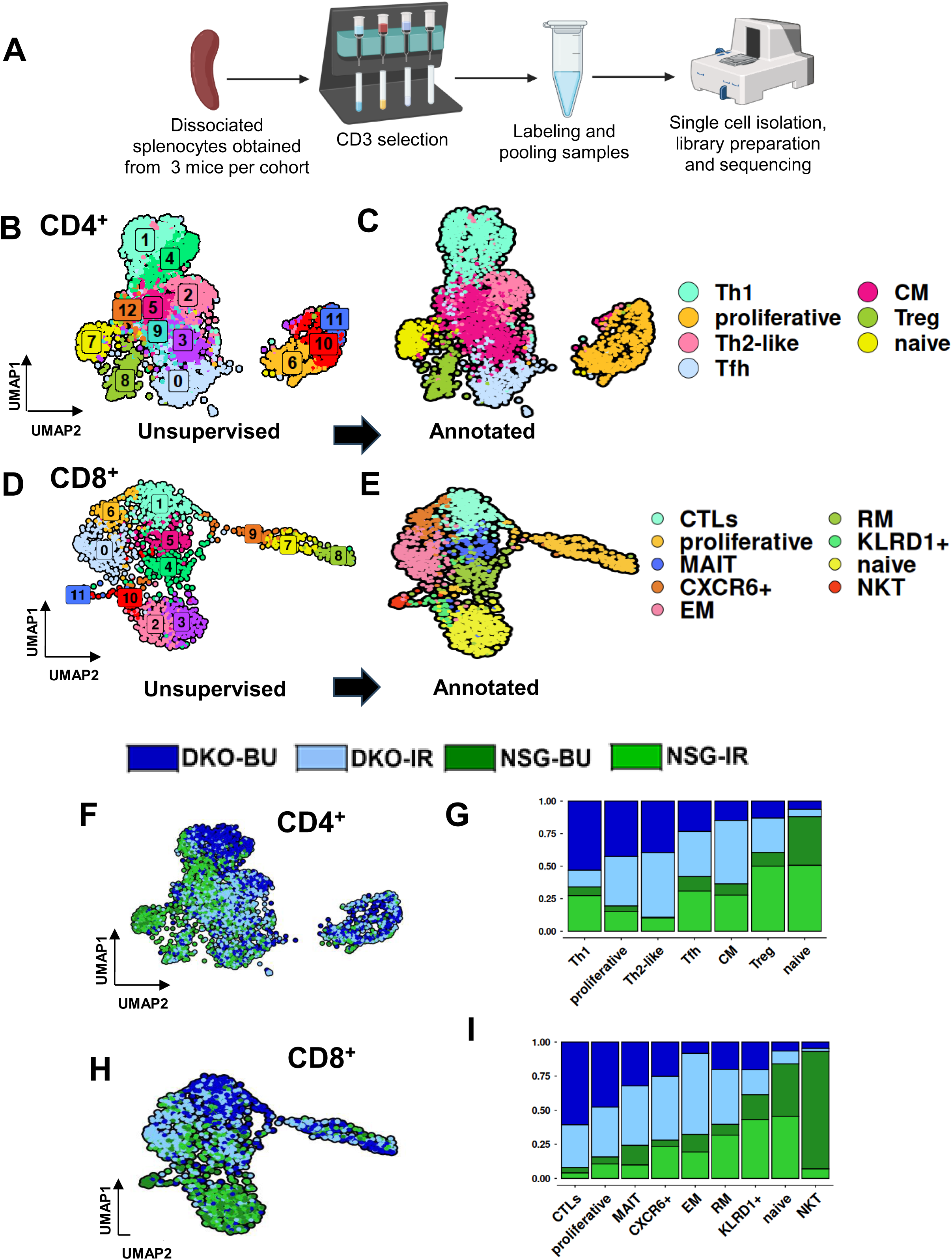
Single-cell RNA-seq analysis of splenic T cells. A. Experimental workflow. Splenocytes were isolated from 3 mice per cohort, dissociated, and subjected to CD3^+^ T cell enrichment. Purified T cells were sample-labeled, pooled, and processed for single-cell isolation, library preparation, and single cell mRNA sequencing. B. Uniform Manifold Approximation and Projection (UMAP) representation of splenic CD4^+^ T cells based on transcriptomic profiling, shown as unsupervised clusters and (C) annotated CD4^+^ T cell subsets. A total of 3,497 CD4^+^ T cells from 12 mice were analyzed and classified. D. UMAP representation of splenic CD8 ^+^ T cells based on transcriptomic profiling, shown as unsupervised clusters and (E) annotated CD8^+^ T cell subsets. A total of 1,644 CD8^+^ T cells from 12 mice were analyzed and classified. F, H. UMAPs of CD4^+^ T cells (F) and CD8^+^ T cells (H) colored according to the experimental cohorts. G, I. Bar plots showing the relative proportions of CD4^+^ T cell subsets (G) and CD8^+^ T cell subsets (I) across the four experimental cohorts.

### Differential gene expression analysis reveals naïve T cell signatures in NSG mice and effector memory signatures in DKO mice

To identify key transcriptional programs distinguishing T cell phenotypes across mouse strains, we performed differential gene expression analysis of CD4^+^ and CD8^+^ T cells. Within the CD4⁺ T cell compartment, comparisons between DKO and NSG mice revealed a clear difference in transcriptional programs (Figure 5A-C). NSG mice, irrespective of conditioning (NSG-BU and NSG-IR), showed significant upregulation of genes associated with naïve and lymphoid-homing T cell patterns, including CCR7, SELL, LEF1, and TCF7. In contrast, these genes were consistently downregulated in DKO mice (DKO-BU and DKO-IR), which instead upregulated activation and effector-associated genes such as TBX21, IFNG, and MKI67 (Figure 5A-C). Notably, T cell factor 7 (TCF7), a transcription factor that maintains T cell stemness and naïve T cell identity, was markedly upregulated in NSG mice while strongly downregulated in DKO mice (Figure 5C). The reduced TCF7 expression in DKO mice is consistent with their elevated effector memory T cell frequencies and more differentiated phenotype. A similar pattern was observed in the CD8⁺ T cell compartment (Figure 5D-F). NSG mice retained high expression of naïve-associated genes (CCR7, SELL, TCF7, LEF1), whereas DKO mice exhibited upregulation of cytotoxic and effector genes, including GZMB, PRF1, GNLY, NKG7, and CCL5, indicative of activated effector memory and cytotoxic T lymphocytes (Figure 5D-F). These transcriptional differences were further accentuated in busulfan-conditioned mice, particularly within the DKO-BU group, which showed the strongest enrichment of cytotoxic effector programs in both CD4⁺ and CD8⁺ T cell compartments (Figure 5B, E). To functionally interpret these gene expression changes, we performed Gene Ontology (GO) enrichment analysis on differentially expressed genes (Figure 5G-J). In DKO mice, upregulated genes in both CD4⁺ and CD8⁺ T cells were enriched for pathways related to cell cycle progression and proliferation (e.g., mitotic nuclear division). Furthermore, CD8⁺ T cells of busulfan-conditioned mice exhibited strong enrichment for cytotoxicity-related pathways, including cell killing and leukocyte-mediated cytotoxicity, further supporting their functional maturation. These findings provide transcriptional evidence that xenogeneic MHC interactions in NSG mice constrain T cell maturation, while their absence in DKO mice permits more physiological differentiation and functional specialization of human T cells.

**Figure 5.**
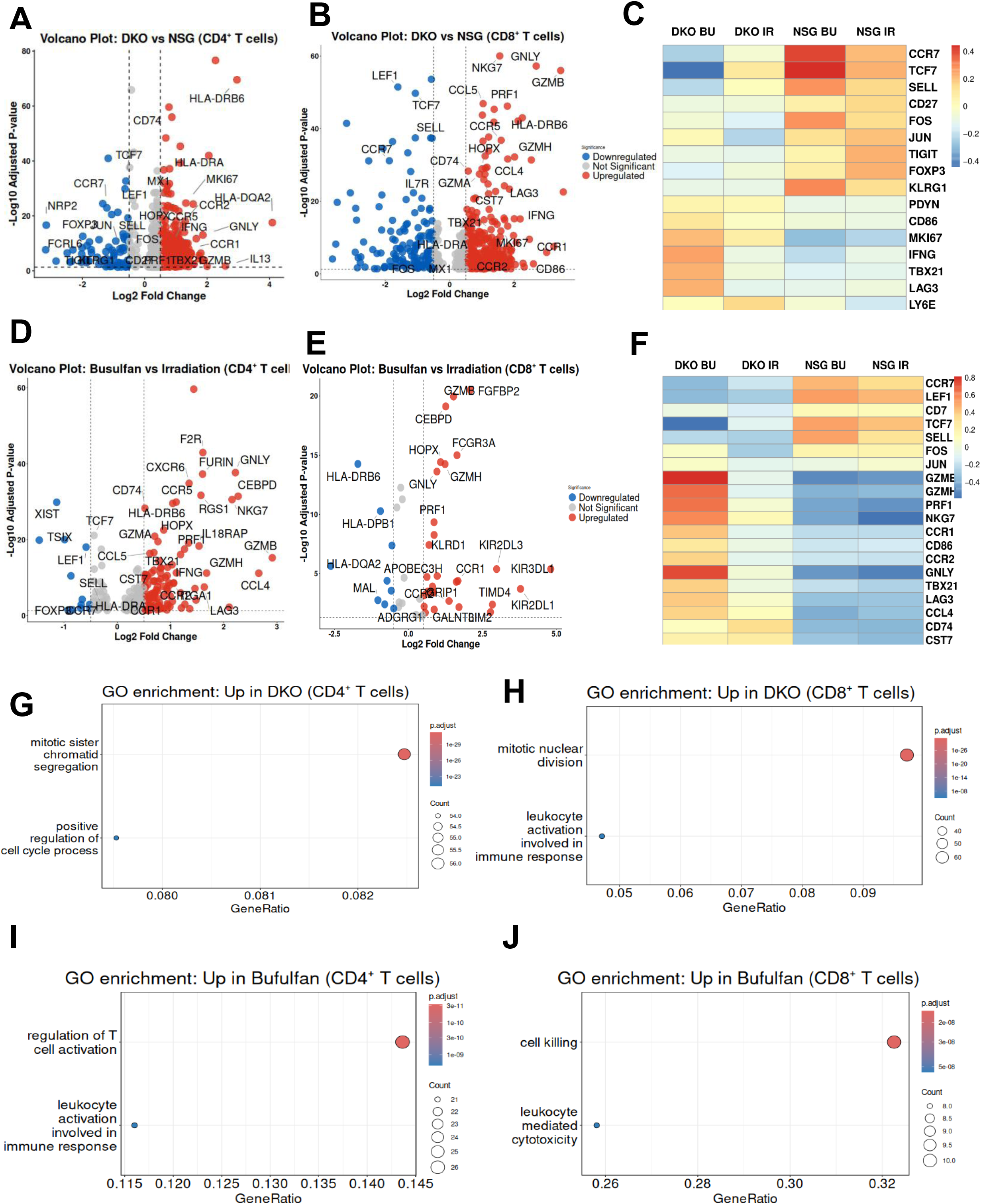
Differential gene expression and pathway enrichment analysis of splenic T cells across experimental conditions. A, D. Volcano plots showing differential gene expression within CD4⁺ T cells, between DKO and NSG cohorts (A) and Busulfan and Irradiation preconditioning (D). The x-axis represents log₂ fold change and the y-axis represents –log₁₀ adjusted p value. Red and blue points indicate significantly upregulated and downregulated genes, respectively. C. Heatmap showing relative expression of selected differentially expressed genes in CD4⁺ T cells across the four experimental cohorts (DKO BU, DKO IR, NSG BU, NSG IR). B, E. Volcano plots showing differential gene expression within CD8⁺ T cells, between DKO and NSG conditions (B) and Busulfan and Irradiation preconditioning (E). The x-axis represents log₂ fold change and the y-axis represents –log₁₀ adjusted p value. Red and blue points indicate significantly upregulated and downregulated genes, respectively. F. Heatmap showing relative expression of selected differentially expressed genes in CD8⁺ T cells across the four experimental cohorts. Colors indicate scaled mean expression levels. G, H. Gene Ontology (GO) enrichment analysis showing the selected biological pathways enriched among genes upregulated in DKO compared with NSG in CD4⁺ T cells (G) and CD8⁺ T cells (H). I, J. GO enrichment analysis showing the selected biological pathways enriched among genes upregulated in Busulfan compared with Irradiation conditions in CD4⁺ T cells (I) and CD8⁺ T cells (J).

### T cells in DKO mice exhibit elevated expression of cytotoxicity genes and activation markers

To further characterize functional T cell phenotypes, we analyzed the expression of cytotoxicity-associated genes, activation markers, and cytokine production within the whole CD3^+^ T cell population. Gene expression analysis revealed significant upregulation of cytotoxic effector genes in DKO-BU mice, including granzyme A (GZMA), perforin 1 (PRF1), and granulysin (GNLY), whereas these genes were downregulated in NSG mice (Figure 6A). To validate these transcriptional findings at the protein level, we quantified human cytokine concentrations in mouse plasma collected at week 20 post-HCT. DKO-BU mice displayed markedly elevated plasma levels of granulysin, perforin, granzyme A, and granzyme B compared to other cohorts (Figure 6B, C). These cytolytic effector molecules are canonical mediators of CD8^+^ T cell cytotoxicity, confirming the functional competence and activated state of CD8^+^ T cells in DKO mice. Analysis of additional cytokines revealed comparable plasma concentrations of interleukin-2 (IL-2) and interferon-gamma (IFN-γ) between DKO and NSG cohorts (Figure 6D). In contrast, interleukin-6 (IL-6) levels were elevated in NSG mice compared to DKO mice (Figure 6E), suggesting distinct inflammatory profiles between the two strains. Collectively, these transcriptional and protein-level analyses demonstrate that T cells developing in DKO mice acquire a highly differentiated cytotoxic phenotype characterized by robust expression of effector molecules. This functional maturation is consistent with the expanded effector memory populations observed in these mice, further supporting the conclusion that the absence of xenogeneic murine MHC permits more physiological human T cell activation and differentiation.

**Figure 6.**
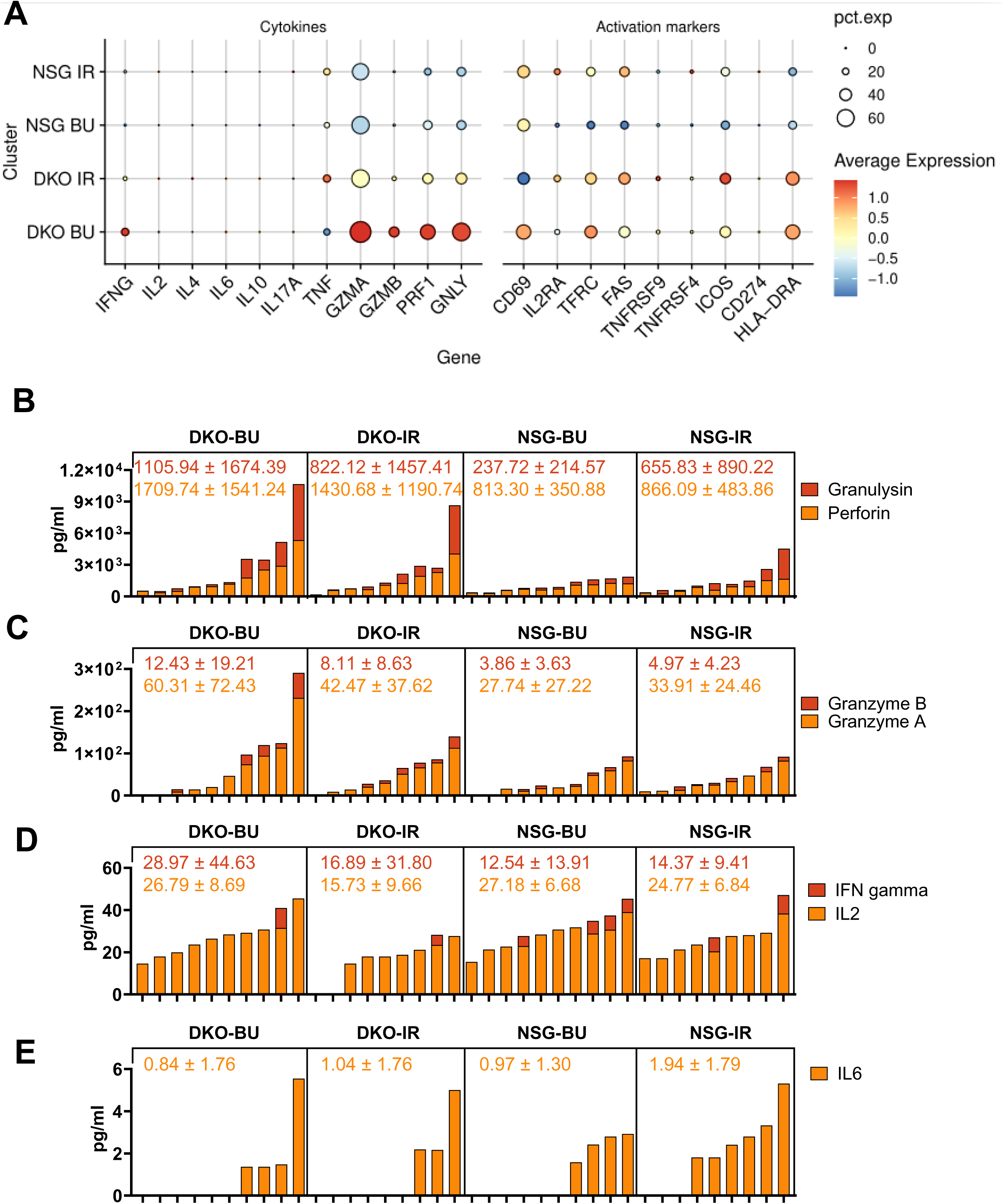
Concordance between transcriptional cytokine signatures in splenic T cells and circulating human cytokines in mouse sera. A. Dot plot showing expression of cytokine and activation marker genes across T-cell populations derived from the four experimental conditions (DKO BU, DKO IR, NSG BU, NSG IR). Dot size represents the fraction of cells expressing the indicated gene, and color intensity indicates the average expression level. B. Serum concentrations of the human cytotoxic proteins granulysin and perforin measured in individual mice across the four experimental cohorts. C. Serum concentrations of the human cytotoxic enzymes granzyme A and granzyme B. D. Serum concentrations of the human inflammatory cytokines IFN-γ and IL-2. E. Serum concentrations of the human inflammatory cytokine IL-6. Bars represent values for individual mice, and numbers above each panel indicate the mean ± s.d. for each cohort. Values below detection threshold are depicted as zero.

### Bulk T cell receptor analyses reveal that DKO mice display a more mature TCR **αβ** repertoire with enhanced clonal expansion

To assess TCR repertoire diversity and clonality, we analyzed TCR sequences from bulk RNA sequencing on CD3^+^ T cells isolated from all mice (n = 10 per cohort) in the DKO-BU and NSG-BU groups, alongside CD34-negative fractions from the original CB-MC units used for humanization. Bulk sequencing analysis revealed significantly higher TCR α and β clonality values in T cells from DKO-BU mice compared to NSG-BU mice (Figure 7A; TCR α clonality: DKO-BU 0.247 ± 0.041, NSG-BU 0.128 ± 0.069; number of unique clonotypes Figure S6A). As anticipated, CB-MC samples exhibited substantially lower clonality values than either mouse cohort, reflecting their higher clonal diversity (Figure 7A and Figure S6A). In contrast, TCR γ and δ chain unique clone numbers were very low in some mice, which could mean that some mice do not seem to develop full TCR γ and δ repertoires (Figure S6A). This trend seemed to be stronger in NSG mice. Analysis of clonal expansion patterns further distinguished the two cohorts. DKO-BU mice harbored the highest frequencies of hyperexpanded TCR α and TCR β clonotypes, followed by NSG-BU mice, while CB-MC samples contained virtually no hyperexpanded clones (Figure 7B). In contrast, no differences in clonal expansion were observed for TCRγ and TCRδ chains between mouse strains, although CB-MC samples exhibited the fewest expanded γδ clonotypes overall (Figure 7C). To further evaluate the functional integrity of the reconstituted human T-cell compartment, we examined CDR3 length distributions across TCR chains as a surrogate measure of repertoire quality. CDR3 length distributions assessed by bulk sequencing were comparable between T cells derived from humanized mice and those from human donors (Figure S6B), indicating that thymic selection and junctional diversification proceed normally in both DKO-BU and NSG-BU models. Single-cell sequencing analyses (Figure S6C) endorsed this conservation of CDR3 length profiles across systems supports the developmental integrity of humanized mouse-derived T cells and validates their use as a model for human T-cell biology. We additionally examined V and J gene segment usage to characterize the repertoire diversification. Humanized mouse-derived T cells demonstrated markedly broader TCR V and J gene utilization compared to CB-MC samples, in which only a restricted subset of genes was detected (examples of TRAJ and TRG V genes can be seen in Figure S7A, B). This expanded gene segment usage in humanized mice likely reflects ongoing thymic output and peripheral T-cell diversification following engraftment, consistent with a maturing immune repertoire. Collectively, these analyses demonstrate that DKO-BU mice, characterized by a predominant effector memory T-cell phenotype, exhibit elevated clonality, restricted TCR diversity, and hyperexpanded clonotypes reminiscent of antigen-experienced adult human T cells — a pattern consistent with the well-established age-associated contraction of TCR diversity driven by memory T-cell accumulation (14). In contrast, NSG-BU mice, which accumulate T cells with a naïve phenotype and less mature transcriptional profiles, maintain broader TCR diversity in line with expectations for antigen-inexperienced T-cell populations (15,16). The enhanced clonality and repertoire restriction observed in DKO-BU mice likely reflect antigen-driven clonal selection and expansion in the absence of xenogeneic murine MHC interference, thereby recapitulating key hallmarks of human adaptive immune maturation.

**Figure 7.**
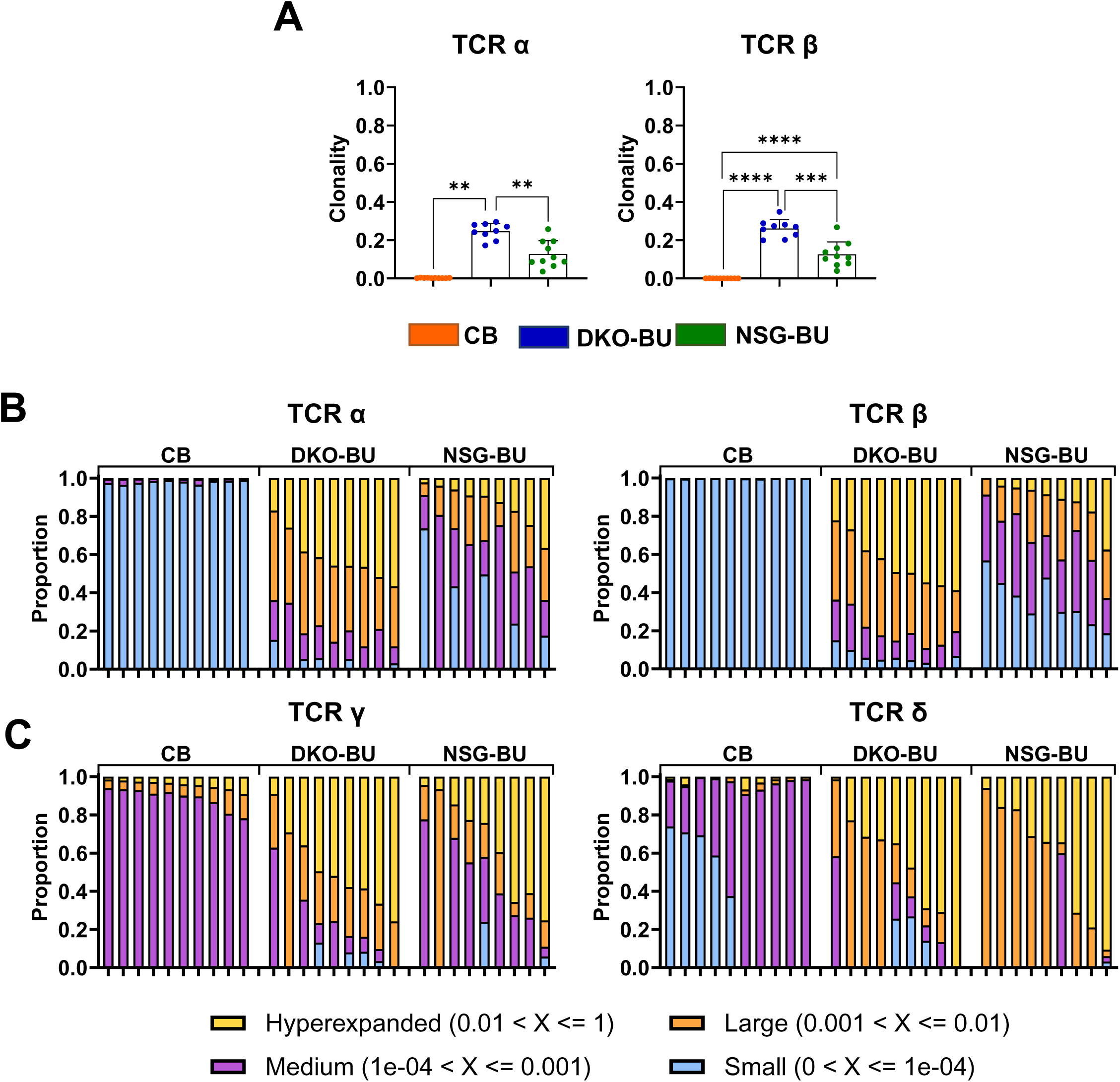
Bulk mRNA sequencing analysis to characterize TCRs expressed by splenic T cells in different cohorts. A. Clonality values obtained from TCRa or TCRb bulk sequencing of CD34neg CB fraction and splenocyte T cells from humanized mice. B. Proportions of TCRa or TCRb populations with different levels of clonal expansion (hyperexpanded, large, medium or small). Bulk sequencing data shown for 9 mice per cohort. CD3+ T cells isolated from 10 cord blood unit (CD34neg CB fraction) were used as reference. C. Proportions of TCRg or TCRd populations with different levels of clonal expansion (hyperexpanded, large, medium or small). Bulk sequencing data shown for 9 mice per cohort. CD3+ T cells isolated from 10 cord blood units (CD34neg CB fraction) were used as reference. DKO-BU (dark blue), DKO-IR (light blue), NSG-BU (dark green) and NSG-IR (light green). Data are presented as mean and SD, and individual values are shown. The differences in A (n = 9 – 10) were analyzed with the Wilcoxon test with Bonferroni correction. **** = p < 0.0001; *** = p < 0.001; ** = p < 0.01; * = p < 0.05. Data are presented as mean and SD, and individual values are shown. The differences in A (n = 9 – 10) were analyzed with the Wilcoxon test with Bonferroni correction. **** = p < 0.0001; *** = p < 0.001; ** = p < 0.01; * = p < 0.05.

## Discussion

In immunocompetent young hosts, T cell development takes place mostly in the thymus, a primary lymphoid organ where bone marrow–derived precursors differentiate into a diverse and self-tolerant T cell repertoire essential for adaptive immunity. Early thymic progenitors enter the thymus (or extrathymic spaces such as liver and spleen) and progress through defined stages, ultimately becoming double-positive (CD4⁺CD8⁺) thymocytes that express a rearranged TCR on their surface (17). Positive selection ensures survival of thymocytes whose TCR can recognize self-peptide–MHC complexes with moderate affinity, while negative selection eliminates strongly self-reactive clones, enforcing central tolerance and preventing autoimmunity (18). Mature single-positive CD4⁺ or CD8⁺ T cells exit the thymus and populate peripheral lymphoid organs as naïve T cells ready to respond to foreign antigens. This coordinated selection process generates a functional, self-restricted T cell repertoire that balances non-self-antigen recognition with self-tolerance.

In humanized mice, human HSPC engraft in the mouse bone marrow, but as the thymus is underdeveloped, T cell precursors must be trained in extrathymic spaces such as the liver and spleen. Over the past decades, significant effort has been devoted into developing mouse models that produce human-specific responses via matching the HLA of HSPC with implanted tissues. Early work by *Brehm et al.* established that humanized NSG mice engrafted with human fetal thymus and liver support the generation of a complete human immune system with HLA-restricted T cells and systemic lymphoid populations (19). Another approach is the bone marrow–liver–thymus (BLT) model, in which human fetal liver and thymus tissues are implanted under the renal capsule of immunodeficient mice, and autologous CD34⁺ HSPC are introduced intravenously. In BLT mice, T cells develop within a human thymic microenvironment, undergoing positive and negative selection on HLA molecules, which yields HLA-restricted T cell repertoires. *Garcia-Beltran et al.* demonstrated that while BLT mice exhibit intact thymic development and TCR diversity, T cell function correlates with levels of innate cells such as monocytes (20). BLT models remain a cornerstone for studying human T cell development, immunopathogenesis, and vaccine or immunotherapy responses *in vivo*; however, the need for human fetal tissues and the technically demanding procedures limit their reproducibility and standardization across laboratories. Significant effort has also been devoted to generating immunodeficient mouse strains in which specific murine MHC molecules are replaced with human HLA alleles. However, due to the extreme polymorphism of HLA molecules, even matching both class I and class II HLA between transgenic mice and the human graft remains technically challenging, limiting the widespread application and reproducibility of these models.

Using a simplified approach, our laboratory previously developed lentiviral (LV)–engineered induced dendritic cells matched to the CD34⁺ HSPC graft, demonstrating that this strategy is sufficient to enhance human T and B cell functional maturation, immune responses against viral antigens, and even promote lymph node regeneration in humanized mice (21–23). However, while robust immune responses to strong viral antigens are commonly observed in these models, eliciting solid responses to weaker tumor antigens remains challenging, as suboptimal TCR development in xenograft systems can mask such reactivity (24).

Building on previous work (12) and confirmed in this study, it is noteworthy that DKO mice can sustain the selection of human single CD4⁺ and CD8⁺ T cells and maintain their long-term reconstitution as activated memory T cells. The distinct CD8⁺/CD4⁺ distribution in blood samples DKO-BU mice may reflect differences in peripheral maintenance, expansion, and tissue distribution rather than selective suppression of CD4⁺ T-cell development, as peripheral blood is a highly dynamic compartment that provides only a snapshot of circulating lymphocytes. Assessment of T cells in lymphoid tissues such as the spleen and bone marrow may therefore provide a more representative measure of overall reconstitution, although direct analysis of trafficking between these compartments remains technically challenging because of the limited blood volume and tissue availability in this model.

Thymic analyses of T cells, thymic epithelial cells, and professional antigen-presenting cells (APC) could have provided important mechanistic insights into T-cell selection in the DKO model. However, these analyses are technically limited in NSG-based humanized mice. Adult NSG mice have a severely hypoplastic thymus as a consequence of the combined *Prkdc*^scid^ mutation and *Il2rγ* deficiency, yielding insufficient tissue for comprehensive flow cytometric and molecular analyses (8). As DKO mice are derived from the NSG background, they share this anatomical limitation. Moreover, mice were analyzed 16–24 weeks after HSC transplantation, when thymic involution was pronounced, and the thymus was frequently barely identifiable macroscopically. These limitations are well recognized in NSG-based humanized mouse models and are one reason why peripheral lymphoid organs are commonly used to assess human immune reconstitution. Although the thymus is the principal site of T-cell development, extrathymic T lymphopoiesis has been reported in the bone marrow, spleen, and secondary lymphoid organs, particularly under conditions of thymic hypoplasia, lymphopenia, or following hematopoietic transplantation (25). Therefore, the relative contributions of thymic selection, extrathymic T-cell development, and peripheral homeostatic proliferation cannot be determined in the present study.

We were also unable to perform an in-depth characterization of human APC because of tissue scarcity. Nevertheless, humanized NSG mice consistently develop abundant human B cells in the spleen, typically comprising more than 30% of human CD45⁺ cells, together with detectable populations of human dendritic cells. These autologous human APC are therefore available to interact with developing and peripheral human T cells. Although the underlying mechanisms remain to be established, we speculate that elimination of murine MHC redirects human T-cell education and activation away from xenogeneic murine MHC recognition toward interactions with autologous human APC. Together with homeostatic signals, these interactions may contribute to the increased frequencies of activated, memory, and cytotoxic T cells observed in DKO mice. This interpretation is consistent with our findings but remains speculative, and future studies will be required to define the relative contributions of extrathymic T-cell development and the role of human antigen-presenting cells in shaping these responses.

It is known that the DKO, compared with NSG or NOG parental strains, shows markedly attenuated GvHD following human PBMC or T cell transfer, with prolonged survival of human T cells detectable in peripheral blood and lymphoid tissues without severe weight loss or xenoreactivity (26). In addition, the transferred human T cells in DKO recipients maintain memory and effector phenotypes and can be used to evaluate immune-modulating therapies over extended periods (26–28). To our knowledge, our current study is the first to demonstrate that the fully humanized DKO mice also show superior T cell development towards mature populations characteristics than the parental NSG mouse strain.

Another important advance in this study is the substantial improvement in the health of DKO mice and their human T cell maturation using myeloablative conditioning with busulfan rather than sublethal irradiation. Total body irradiation damages mitotically active cells and is efficient for the reduction in lymphocyte populations in order to generate niche space within the bone marrow compartment of recipient mice for HSPC engraftment. Busulfan is a myeloablative alkylating agent and functions in depleting non-cycling primitive stem cells (29). Several lines of evidence in NOD/SCID and NSG mice showed that busulfan conditioning supports efficient human hematopoietic stem cell engraftment with comparable or improved reconstitution while avoiding some of the toxicity and logistical challenges associated with irradiation. Pre-conditioning with busulfan enables levels of human CD45⁺ chimerism that are similar to or exceed those achieved with sublethal irradiation, with improved survival and lower animal stress, and supports multilineage reconstitution, including B and T lineage cells (30,31). The effects of myeloablative busulfan conditioning on human myeloid-cell development were not directly investigated in this study. However, busulfan conditioning is generally considered to induce less acute tissue injury and inflammatory signaling than total-body irradiation while efficiently creating hematopoietic niche space for donor HSC engraftment. Consequently, busulfan-conditioned mice may provide a less inflammatory environment for immune reconstitution, potentially favoring physiological differentiation of human myeloid cells and their interactions with developing T cells. Future studies should define how different conditioning regimens influence human myeloid-cell development and function in next-generation humanized mouse models. Beyond its biological effects, the conditioning regimen also has important practical implications for the development and broader adoption of humanized mouse models in preclinical cell and gene therapy research. Busulfan conditioning represents an attractive alternative to total-body irradiation because it can be administered without specialized irradiation facilities, which are not readily available at many research institutions. In addition, irradiation protocols require careful dosimetric calibration and standardization between facilities to ensure reproducible hematopoietic ablation while minimizing toxicity. In contrast, busulfan provides a more accessible and readily standardized pharmacological conditioning approach that facilitates implementation across laboratories. The enhanced human T-cell development observed in busulfan-conditioned DKO mice further supports the use of this conditioning strategy for preclinical studies evaluating advanced cell and gene therapies and requires further investigation.

Giusti et al. documented the progressive development of granulomatous lesions. Giusti and colleagues reported spontaneous granulomatous lesions in humanized NSG mice, with an incidence of approximately 80% by 26 weeks and 90% by 35 weeks post-engraftment, most frequently affecting the spleen, liver, and bone marrow and occasionally containing multinucleated giant cells (32). In our study, humanized NSG mice analyzed at approximately 20 weeks post-HCT predominantly exhibited small lymphoid cells, without the prominent large-cell morphology characteristic of the granulomatous pathology described by Giusti et al. Some DKO mice, however, showed more prominent clusters of F4/80⁺ murine myeloid cells surrounding human CD4⁺ large-cell aggregates. Determining whether these findings represent a relevant tissue phenotype would require longitudinal analysis of mice maintained to later time points together with comprehensive characterization of both human and murine myeloid compartments. A further limitation of the present study is that the identity of the enlarged human CD4⁺ cells detected by immunohistochemistry could not be conclusively established. Although their morphology was initially interpreted as consistent with activated CD4⁺ T cells, human CD4 is also expressed by subsets of monocytes, macrophages, and dendritic cells. Additional immunohistochemical analyses to detect human CD68 were inconclusive and therefore did not permit reliable discrimination between human CD4^+^ lymphoid and myeloid populations. Such analyses were beyond the scope of the present study and could not be performed because the limited tissue available was prioritized for detailed T-cell analyses by flow cytometry and RNA sequencing. Nevertheless, systematic characterization of murine and human myeloid development and late-stage immune infiltrations in the spleen of DKO mice represents an important direction for future studies.

The low or undetectable numbers of individual TCR clones observed in some animals should be interpreted with caution, particularly for the single-cell TCR dataset, because the number of cells available for this analysis was limited and not every captured cell yielded a productive TCR sequence. We therefore complemented the single-cell analysis with bulk TCR sequencing to increase the number of analyzed T cells and enable more robust quantitative comparisons between experimental groups. The predominance of αβ over γδ TCRs is consistent with the distribution of T-cell subsets in humans, in which αβ T cells constitute the major circulating T-cell population. Therefore, the lower representation of γδ TCRs in humanized mice was expected. While altered T-cell selection and peripheral expansion may contribute to the observed repertoire differences, the present data do not allow us to determine whether the DKO-BU environment specifically alters thymic αβ TCR selection or actively directs progenitor differentiation toward a particular TCR pathway. Such mechanistic analyses would require dedicated investigation of TCR rearrangements at earlier stages of human T-cell differentiation.

The single-cell and bulk TCR analyses provide complementary but distinct information on T-cell clonality. Single-cell RNA/TCR sequencing permits direct assignment of individual TCR clonotypes to their transcriptional phenotype, whereas bulk TCR sequencing provides greater depth and sensitivity for quantifying expanded clonotypes but does not retain the cellular transcriptional phenotype of individual clones. Higher frequency of hyperexpanded clonotypes detected by bulk TCR sequencing in DKO-BU mice reflects the greater depth of this approach and cannot be directly assigned to individual transcriptomic clusters. This complementary analysis supports the presence of expanded T-cell populations with activated, cytotoxic, and proliferative phenotypes in DKO-BU mice while avoiding overinterpretation of the bulk repertoire data.

The distinct inflammatory profiles of NSG and DKO mice may reflect differences in xenoreactive stimulation. In NSG mice, human T cells can recognize murine MHC molecules, potentially promoting persistent low-grade T-cell and myeloid activation and systemic inflammatory signaling, including increased IL-6, consistent with features of xGVHD- or CRS-like inflammation. In contrast, deletion of murine MHC in DKO mice reduces this xenoreactive stimulus and may permit greater cytokine-supported homeostatic expansion and differentiation of human T cells following busulfan-mediated lymphodepletion, resulting in enhanced cytotoxic programs, including Granulysin, Perforin and Granzymes, without increased systemic IL-6. The apparently discordant IFNG findings should also be interpreted in the context of the different analytical compartments: IFNG transcripts were readily detected in splenic T cells by scRNA-seq and were significantly increased in DKO mice, whereas circulating IFN-γ protein was below the detection threshold in many plasma samples. Thus, the plasma measurements do not provide sufficient quantitative resolution to determine differences in systemic IFN-γ concentrations, and tissue-level IFNG transcription should not be expected to translate directly into detectable circulating protein, particularly when cytokine production is localized to activated T-cell populations within lymphoid tissues.

A limitation of the present study is that the functional activity of the reconstituted human T cells was not directly assessed in antigen-specific *ex vivo* or *in vivo* challenge assays. The increased expression of cytotoxic effector molecules and activation-associated programs therefore demonstrates cytotoxic potential rather than definitive antigen-specific effector function. Because this study was designed as a baseline characterization without tumor, infection, or immunization and tissue availability was limited, functional challenge experiments were beyond its scope. Future studies should determine the functional activity and antigen specificity of both cytotoxic and helper T-cell populations against defined cancer or infectious targets.

In the present study, only female DKO mice were humanized, as female recipients have been reported to support higher levels of HSPC engraftment and more efficient T cell development compared with males (33,34). It is not known, therefore, whether male DKO mice would show the same effects observed here for females. By the way, researchers developed the THX humanized mouse model by transplanting human CD34⁺ cells and then conditioning mice with 17β-estradiol. Estradiol treatment promoted differentiation of human lymphoid and myeloid cells, including T follicular helper cells and germinal center B cells with mature antibody responses (35). Similar studies remain to be conducted with humanized DKO mice.

In summary, DKO-derived humanized mouse models offer a versatile platform for studying human immune development and function, but their potential can be further enhanced through targeted cytokine knock-ins. Expression of human IL-7 significantly improves T cell differentiation, survival, and peripheral reconstitution in humanized mice, while provision of human FLT3L promotes the development of diverse human dendritic cell subsets capable of autologous HLA-matched antigen presentation (36,37). Notably, models incorporating FLT3L signaling, such as the Flt3-deficient humanized mice described by Di Santo and colleagues (38), demonstrate robust reconstitution of human antigen-presenting cells and innate immune compartments. Collectively, these cytokine-refined humanized models provide a more physiologically relevant platform for investigating human adaptive and innate immunity, vaccine responses, and immunotherapeutic strategies, and set the stage for the next generation of HLA-matched precision humanized mouse models.

## Materials and methods

### Cord blood selection

The study was conducted according to the guidelines of the Declaration of Helsinki. The collection of CB specimens was approved by the Ethics Committee of the Hannover Medical School (MHH) and was performed by the Research Obstetric Biobank (approval number 1303 to Constantin von Kaisenberg). The use of the CB specimens to humanize mice was approved by the Ethics Committee of the MHH (approval number 4837 to Renata Stripecke). The transfer of the pseudo-anonymized CB samples from the MHH to the Stripecke lab for use at the University Hospital of Cologne (UKK) was approved by the UKK Ethics Committee (approval number 22-1423_3 to Renata Stripecke). All cord blood products received an anonymous numerical code, by which no donor could be identified. After CB collection, MC CD34^+^ cells were positively selected through two consecutive runs using an immune magnetic bead kit (Miltenyi Biotec, Bergisch Gladbach, Germany) and cryopreserved as previously described (21,39). The HLA genotypes of ten CB units were obtained through sequencing using reported methods (Table S1) (12). The generated sequences were analyzed using NGSengine version 3.2.0, and the IMGT 3.57.0 database was implemented.

### Generation of humanized mice

The animal protocols for mouse studies were approved by the animal care committee of the state of North Rhine-Westphalia (LAVE, approval number Az 81-02.04.2023.A154 to Renata Stripecke) and performed according to the German animal welfare act and the EU directive 2010/63. All animals were housed in individually ventilated cages in a specific and opportunistic pathogen-free environment. The well-being of mice was monitored according to score sheets approved by LAVE with pre-defined humane endpoints (Table S2). The parental NSG (stock number 005557, *NOD.Cg-Prkdc^scid^ IL2rg^tm1Wjl^/SzJ*) and derived DKO (stock number 025216, *NOD.Cg-Prkdc^scid^ H2-K1^b-tm1Bpe^ H2-Ab1^g7-em1Mvw^ H2-D1^b-tm1Bpe^ IL2rg^tm1Wjl^/SzJ*) mice (females, 5 weeks-old) were obtained from The Jackson Laboratory (Bar Harbor, MA, USA) and housed in our facility for a week prior to transplantation. For preconditioning, the mice were either irradiated (150 cGy) using an X-ray irradiator (MultiRad 160, Faxitron, Tucson, Arizona, USA) 4 h prior to HCT or injected intraperitoneally with 100 µl of 15 mg/kg Busulfan (Fresenius Kabi, Bad Homburg vor der Höhe, Germany) 24 h prior to HCT according to methods described in a STAR protocol (40). CD34^+^ CB-MC obtained from ten different donors were chosen for HCT based on the number of viable CD34^+^ cells and low frequency of remaining T cells (less than 3%). The animals received antibiotics (Cotrim-K, Ratiopharm, Ulm, Germany) in their water two days before preconditioning and continued receiving them during the experiment. 2 x 10^5^ viable human CB CD34^+^ cells were injected i.v. into the tail vein of mice.

### Acquisition of reference human peripheral blood mononuclear cells

PBMC were obtained from healthy blood donors. The samples were pseudonymized and kindly provided by the University of Cologne Blood bank following informed consent (Ethics protocol number 23-1309). The PBMC were purified by density-gradient centrifugation with Pancoll (PAN Biotech, Aidenbach, Germany). The buffy coats were harvested, washed, and cryopreserved as viable cells at -150°C.

### Blood and plasma collection

Human immune reconstitution in the peripheral blood lymphocytes was monitored at 8, 12, 16, and 20 weeks post-HCT. Around 100 µl of blood was collected by submental bleed at weeks 8, 12, and 16. The red blood cells were removed by adding 100 µl red blood cell lysis buffer (RBC, Invitrogen, Waltham, Massachusetts, USA) and incubating for 10 min at room temperature. Then, the reaction was stopped with 100 µl FACS buffer (PBS + 1% FCS), and samples were centrifuged for 5 min at 300 x g. If the pellets showed observable hemoglobin, the procedure was repeated. At 20 weeks post-HCT, terminal blood was collected by cardiac puncture after sacrificing the animals. Around 100 µl of blood was taken for flow cytometry analysis and processed as described before (12). The rest was centrifuged for 20 min at 2500 x g, and plasma samples were collected for further analysis.

### Multiplex cytokine analysis

Cytokine concentrations in the mouse plasma were determined using the LEDGENDplex^TM^ Human CD8/NK Panel Multi-Analyte Flow Assay (BioLegend, San Diego, California, USA), which allowed for the simultaneous detection of human interleukin 2, 6, granulysin, granzyme A, granzyme B, interferon (IFN)-γ, and perforin. The assay was performed according to the manufacturer’s instructions and using ¼ of all reagents. Samples were analyzed with the Data Analysis Software Suite for LEGENDplex™ version 2024-06-15 (BioLegend, San Diego, California, USA). The values not reaching the minimum detectable value for each cytokine according to the assay were depicted as 0 values.

### Tissue collection and immunohistochemistry

Pieces of spleen, heart, lung, kidney, liver, gut, and thymus were fixed in 10 % formalin solution (Sigma Aldrich, St. Louis, Missouri, USA) overnight. Following a fixation period of 24 h, the spleen was embedded in paraffin. Sections were cut at three micrometers, dewaxed, and subjected to immunohistochemistry staining as described (41). Briefly, heat-induced antigen retrieval was initially performed in a citrate buffered solution. All slides were rinsed with Tris-buffered saline (pH 7.6) plus 0.01% Tween R 20 (Merck, Darmstadt, Germany). Slides were incubated for 20 min at 21°C in normal goat serum (Vector Laboratories Inc., Burlingame, CA) and then incubated with the primary antibody for 1 h at 21°C (anti-human nuclei or anti-CD4). Secondary biotin-SP-conjugated antibody was applied for 30 min at 21°C. Final staining was achieved by the routine method using alkaline phosphatase streptavidin-biotin (Dako REAL-Strep AP, Agilent, Santa Clara, CA, USA) and red as chromogen (Dako REAL-chromogen red, Agilent). The slides were counterstained using Bluing Reagent (Thermo Scientific, Braunschweig, Germany). To characterize macrophages within the spleen, primary antibodies directed against human macrophages CD68 or mouse macrophages F4/80 were used. Pre-treatment was performed with protease (CD68 staining) or proteinase k (F4/80 staining) followed by incubation with the primary antibody for 1 h at 21◦C (CD68) or overnight at 4 °C (F4/80). Staining was achieved by routine method using alkaline phosphatase streptavidin-biotin (Vector Laboratories, Newark, California, USA) and DAB (3, 3’-diaminobenzidine) as chromogen (Carl Roth GmbH + Co. KG, Karlsruhe, Germany). Sample permeabilization, antibody concentrations, antibody reactions, and staining procedures were previously optimized for each antibody (41) to get clear and specific immunohistochemical signals. Slides were digitized using the Hamamatsu S-210 Nanozoomer scanner (Hamamatsu, Herrsching am Ammersee, Germany) equipped with a 20x plan-apochromat lens. Information about the antibodies used can be found in Table S3.

### Tissue collection, processing, and preservation

The remaining spleen and bone marrow samples were prepared as single-cell suspensions. Red blood cells were removed by incubation with RBC lysis buffer for 10 min at room temperature. The total viable cells were counted, 5 x 10^6^ cells were used directly for flow cytometry analysis, and the remainder were either used for T cell isolation (spleen) or cryopreserved as viable cells at -150°C.

### Blood and tissue analysis by flow cytometry

Freshly isolated single cell suspensions of were processed with mouse tissues (blood, spleen, and bone marrow) were blocked for 15 min at 4°C with mouse IgG block (Sigma Aldrich, St. Louis, Missouri, USA). For human cryopreserved CB and PBMC reference samples, cells were thawed and analyzed. For exclusion of dead cells during analyses, cells were stained with Zombie UV dye (Biolegend, San Diego, California, USA). Afterwards, cells were stained with monoclonal antibodies for flow cytometry for 20 min at 4°C (Table S3). Data were acquired on a Cytoflex LX flow cytometer (Beckman Coulter, Brea, California, USA) and analyzed using the Kaluza software version 2.3 (Beckman Coulter). Gating strategies can be seen in Figures S8 and S9.

### T cell isolation

T cells were isolated from single-cell suspensions of spleens using CD3 microbeads. In detail, the cells were labeled using CD3 microbeads according to the manufacturer’s instructions and applied to the prepared MACS LS columns (Miltenyi Biotec) attached to the magnet. Firstly, the negative fraction was collected by washing the columns 3 times using 3 ml of rinsing buffer (Miltenyi Biotec). Then, the columns were removed from the magnet, and the positive fraction was collected by applying 5 ml rinsing buffer and plunging them. Both negative and positive fractions were centrifuged, and the supernatants were removed. The cells were resuspended and counted. The cells were either used for single-cell mRNA analysis (as described below) or frozen at -80 °C as dry pellets, which were later used for bulk mRNA analysis.

### Single-cell mRNA sequencing (scRNA-seq) analysis

Single-cell RNA sequencing was performed on isolated T cells from spleen samples using the BD Rhapsody™ Express Single-Cell Analysis System. Sample multiplexing was achieved using the BD Mouse Single-Cell Multiplexing Kit (BD Biosciences, San Jose, CA, USA). Briefly, up to 1 × 10 cells per sample were incubated with 10 µl of oligonucleotide-conjugated hashtag antibodies at room temperature for 20 min, washed twice to remove unbound oligonucleotides, counted, and pooled at equal representation. Single cells were isolated via the Single-Cell Capture and cDNA Synthesis with the BD Rhapsody Express Single-Cell Analysis System according to the manufacturer’s recommendations (BD Biosciences) using the BD Rhapsody Cartridge Kit (BD Biosciences) and the BD Rhapsody cDNA Kit (BD Biosciences). Whole transcriptome, sample tag, and TCR/BCR libraries were prepared using the BD Rhapsody Whole Transcriptome Analysis (WTA) Amplification Kit (BD Biosciences) and the BD Rhapsody TCR/BCR Amplification Kit (BD Biosciences) following the BD Rhapsody System TCR/BCR full length, mRNA WTA and Sample Tag Library Preparation Protocol (BD Biosciences). Final libraries were quantified using a Qubit Fluorometer with the Qubit dsDNA HS Kit (Thermo Fisher Scientific, Waltham, MA, USA), and fragment size distributions were assessed using the Agilent high-sensitivity D5000 assay on a TapeStation 4200 system (Agilent). Libraries were sequenced on an Illumina NovaSeq X platform in paired mode with a read configuration of 86 bp for read 1 and 216 bp for read 2, using NovaSeq X 10B Reagent Kit (300 cycles) chemistry. After demultiplexing of bcl files using Bcl2fastq2 V2.20 (Illumina) to generate fastq files the BD Rhapsody Sequence Analysis Pipeline 2.2 was used to align reads against the gencode human reference genome (hg38) and generate molecular counts per bioproduct per identified nucleus and metrics related to the different steps (42). Alignment was conducted using the RhapRef_Human_WTA_2023-02 reference. scRNA-seq count matrices were processed and analyzed in R (v4.5.0) using the Seurat package (v5.3.0) (43). Hashtag counts were normalized. Cells identified as singlets were retained for downstream analyses. Low-quality cells were excluded based on the number of detected genes and the percentage of mitochondrial genes (specific thresholds for samples obtained from each mouse). Importantly, ribosomal genes (small and large subunits) as well as mitochondrial genes with MT-identifier were excluded from the analysis. The normalization method was set to ‘LogNormalize’. To delineate CD4^+^ T cells, the object was subsetted based on normalized expression with the following thresholds: CD4 expression greater than 0.1, CD8A expression less than 0.1, and TRDC expression less than 1. For CD8^+^ T cells, the object was subsetted based on the following thresholds: CD8A expression greater than 0.1, CD4 expression less than 0.1, and TRDC expression less than 1. Dimensionality reduction was performed using the RunUMAP function with dimensions 1:30. The default resolution was used for clustering. To characterize the clusters, differential gene expression analysis was performed using the FindMarkers function in Seurat. TCR repertoire data were processed, analyzed, and integrated with scRNA-seq data using the scRepertoire package (v2.5.3) (44). While performing the TCR repertoire analysis using scRepertoire, the cloneCall parameter was set to ‘strict’, which uses the V(D)JC genes comprising the TCR plus the nucleotide sequence of the CDR3 region to call the clonotypes. Both chains were used for the clonotype analysis wherever detected. Shannon’s diversity score was calculated based on amino acid sequences of the TCR repertoire. Gene Ontology (GO) pathway analysis of the differentially expressed genes was performed using the clusterProfiler R package (version 4.18.4) (45).

### Bulk mRNA sequencing analysis

The frozen pellets of isolated T cells from spleen samples were used for RNA isolation using QIAGEN AllPrep DNA/RNA Mini-Kit (Qiagen, Hilden, Germany) according to the manufacturer’s instructions. RNA integrity was determined using Agilent TapeStation4150, with High Sensitivity RNA Screen Tapes (Agilent). RNA-based TCRβ-chain (TRB) deep-sequencing was performed using DriverMap-AIR TCR Profiling-Kit (Cellecta, Mountain View, California, USA) according to the manufacturer’s protocol, using 200 ng (CB-MC) or 200-900 ng (spleen, depending on yield) RNA as input. RNA-based TRB libraries were sequenced on Illumina NovaSeq6k (300 cycles) by the DKFZ Next Generation Sequencing Core Facility. Alignment of raw sequencing output was performed using mixcr with cellecta-human-rna-xcr-umi-drivermap-air preset (46). Data alignment, analysis, and visualization were performed using mixcr (v4.7.0) (46), VDJtools (v1.2.1) (47), and R-based tools (immunarch v0.6.6; ggplot2) (48). Diversity is the total number of unique clonotypes in a sample. TCR alpha and beta diversity were downsampled for quantitative reasons. Clonality is defined as ‘1 – Shannon’s entropy/log2(diversity)’ (49).

### Statistics and data visualization

Statistical analysis was carried out with the statistical software R version 4.5.0. All mean values with standard deviations and calculated p-values can be found in Tables S5-S9. Visualization of gene expression and single-cell TCR data was carried out using ggplot2 (version 3.5.1), Scpubr (version 2.0.2) (50). The visualization of other data was done using GraphPad Prism software version 10.4.1 or with R’s heatmap2 function, chain usage is annotated using VDJtools

## Data availability

The single-cell sequencing data can be found in the BioStudies database (http://www.ebi.ac.uk/biostudies) under accession E-MTAB-17275.

## Acknowledgements

This work was sponsored by The Jackson Laboratory (Grant LV-HLA, LV-HLA2) as an academic collaboration contracted grant (to RS). Development of humanized mouse models at the Stripecke laboratory is also funded by the Cancer Research Center Cologne Essen (CCCE), German Cancer Aid (Deutsche Krebshilfe GRANT 70116565), and Hector Foundation (GRANT M2418) (to RS). This work was also supported by NIH grant UG3DK142192 to LDS. ZZ is supported by the Deutsche Forschungsgemeinschaft (DFG, German Research Foundation) – 322977937/GRK2344. MDB is a member of the excellence cluster ImmunoSensation3 (EXC2151 project number 390873048). This work was supported by the Ministry of Culture and Science of North Rhine-Westphalia. NGS analyses were carried out at the Joint Scientific Facility WGGC-Bonn PRECISE.

## Author Contributions

MD planned experiments, acquired mice and reagents, performed mouse monitoring and collection and processing of tissues for analyses, procured data, provided samples to collaborators, performed analyses with collaborators, prepared the figures and tables, wrote the draft of the manuscript, and performed revisions. ZZ performed biocomputational single cell sequencing analysis and prepared transcriptomic analysis figures. FK performed the i.v. injections of HSPC and assisted in mouse monitoring, collection, and processing of tissues for analyses. MR tested the cryopreserved HSPC batches and assisted in mouse monitoring and collection, and processing of tissues for analyses. S supervised bioinformatic analyses of single cell sequencing data. AE and IP performed bulk TCR sequencing and data preparation. JS-S, ED-D, and MDB performed single cell sequencing and data preparation. DS performed immunohistochemistry analysis of embedded tissues. AD and BE-V performed cytokine analysis of the mice’s plasma. CvK procured and provided CB units. FK performed statistical analysis of all data. MT and HAS provided optimized pretested panels for multicolor FACS analyses in Cytoflex. EB performed HLA genotyping of CB. FK and AS assisted with the operation of the irradiation device. LDS and BS provided conceptualization of studies to test DKO mice, knowledge to present the data, and revised the manuscript. RS planned the project, designed experiments, acquired funding, supervised the project, supervised data analyses and preparation of figures, wrote parts of the manuscript, and edited the final versions. All authors reviewed and approved the manuscript.

## Conflict of Interest

R.S. obtained research support and received honoraria for participating in and organizing conferences with The Jackson Laboratory (JAX), a not-for-profit organization commercializing the mouse strains used in these studies. MD participated in conferences sponsored by JAX. L.S. and B.S. are employees of JAX. Other authors declare no conflict of interest.

