## Supplemental Figures for "Memory T Cells in MHC-Deficient Humanized Mice"

Figure S1: Analyses of weight and human T cell frequencies.

A. Weights of individual mice per each cohort (in grams). One DKO-IR and one NSG-IR mouse succumbed during the experiment.

B. Analyses of $CD3^{+}$ cell fraction within huCD45 in blood at weeks 8, 12, 16, and 20 after HCT (in percentages).

C. Analyses of $CD4^{+}$ cell fraction within CD3/CD45 in blood at weeks 12, 16, and 20 after HCT (in percentages).

D. Analyses of $CD4^{+}$ and $CD8^{+}$ cells in bone marrow (in percentages)

E. Analyses of $CD4^{+}$ and $CD8^{+}$ cells in spleen (in percentages).

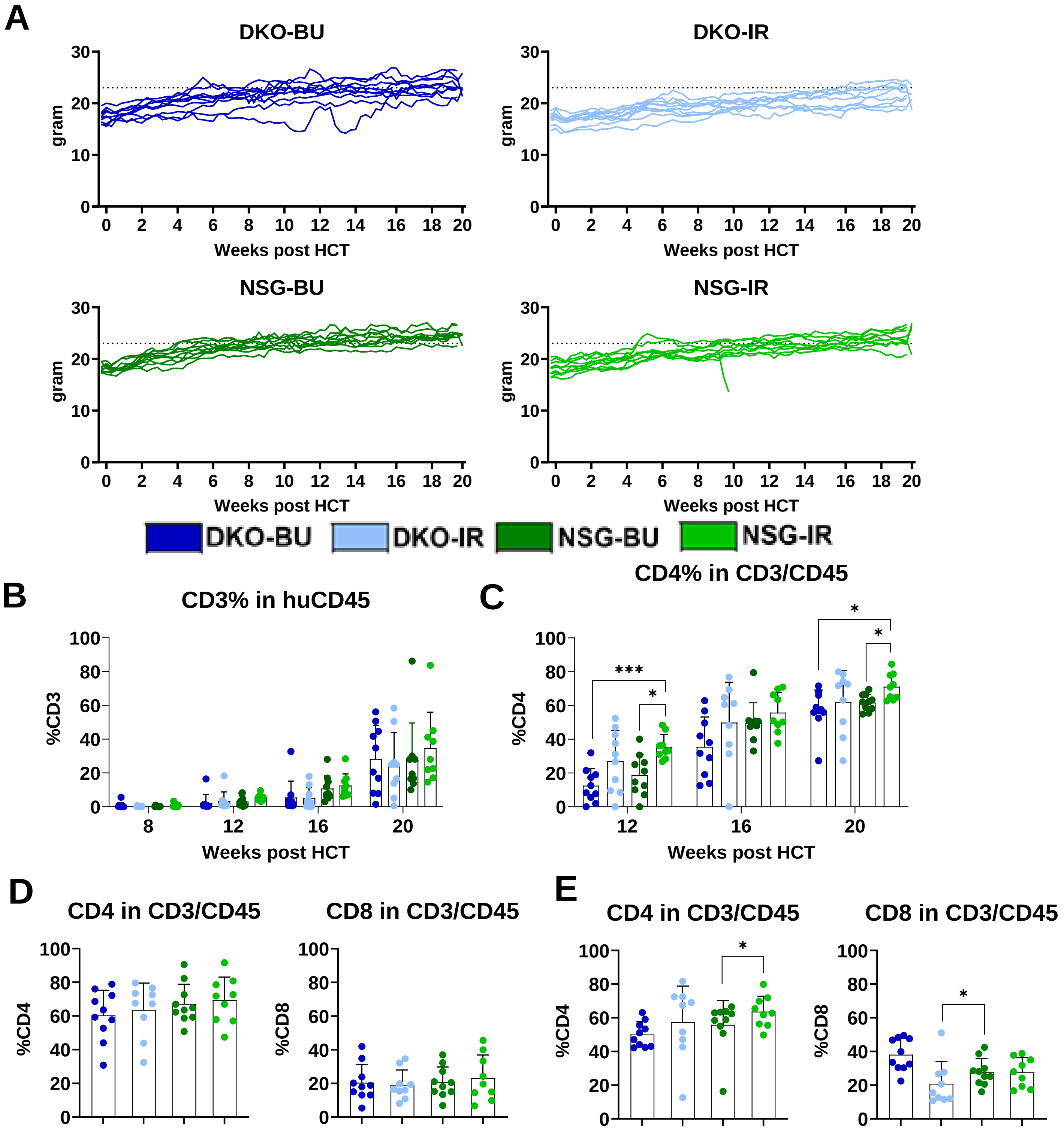

Figure S2: T cell phenotype in adult PBMC (magenta), $CD34^-$ CB-MC (orange), and tissues of humanized mice (spleen /SPL, peripheral blood/ PBL, and bone marrow/ BM). DKO-BU (dark blue), DKO-IR (light blue), NSG-BU (dark green), and NSG-IR (light green).

A. Frequencies of central memory T cells within the $CD4^+$ fraction (in percentage).

B. Frequencies of terminal effector T cells within the $CD4^+$ fraction (in percentage).

C. Frequencies of central memory T cells within the $CD8^+$ fraction (in percentage).

D. Frequencies of terminal effector T cells within the $CD8^+$ fraction (in percentage).

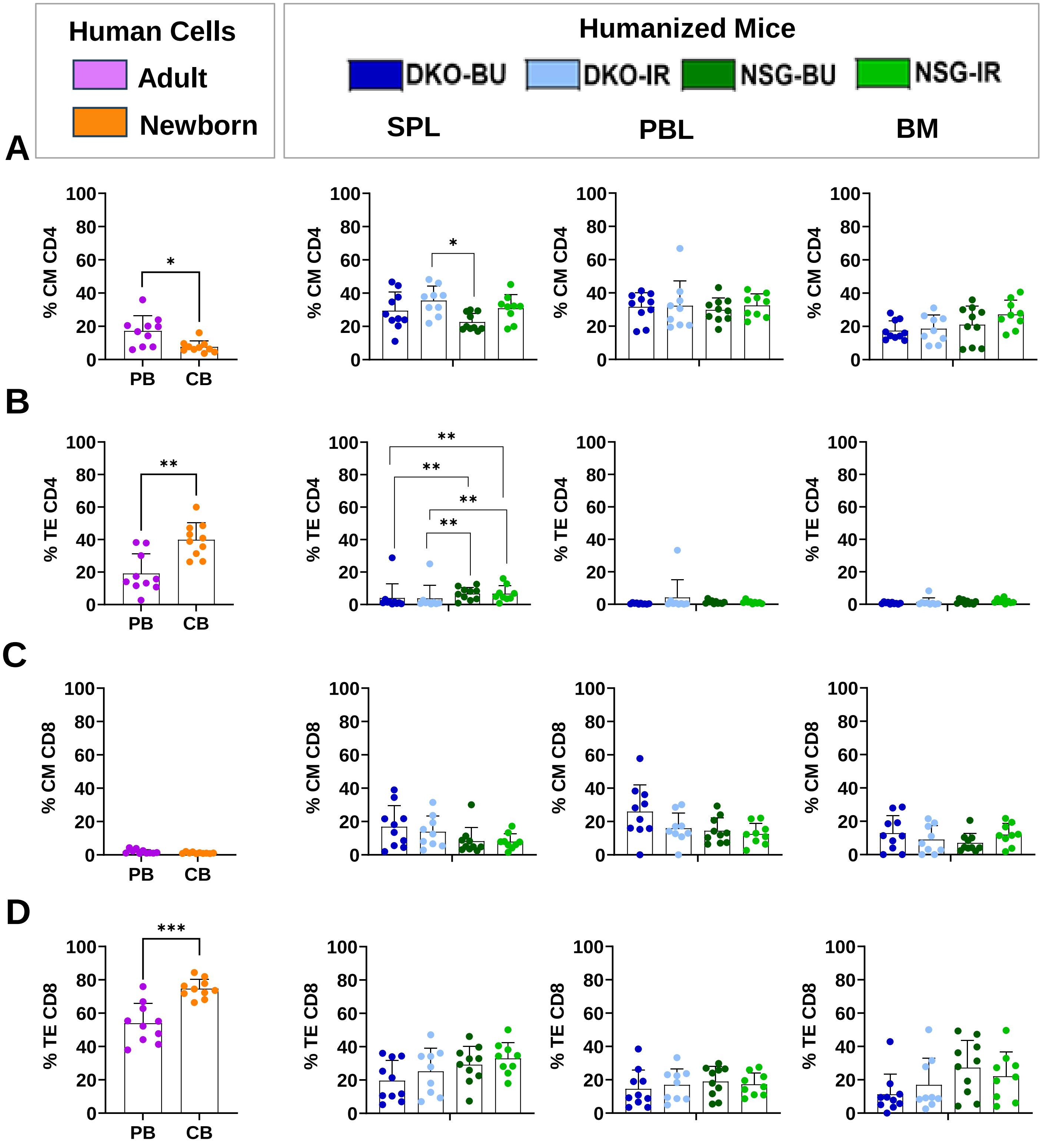

Figure S3: Total T cell counts in spleen for each mouse grouped per CB unit used for humanization. DKO-BU (dark blue), DKO-IR (light blue), NSG-BU (dark green) and NSG-IR (light green) (in counts). The mice used for single-cell mRNA sequencing for cord blood units 74, 160 and 161 are marked by red squares. X marks mice that were lost in the course of the experiment.

Individual values are shown.

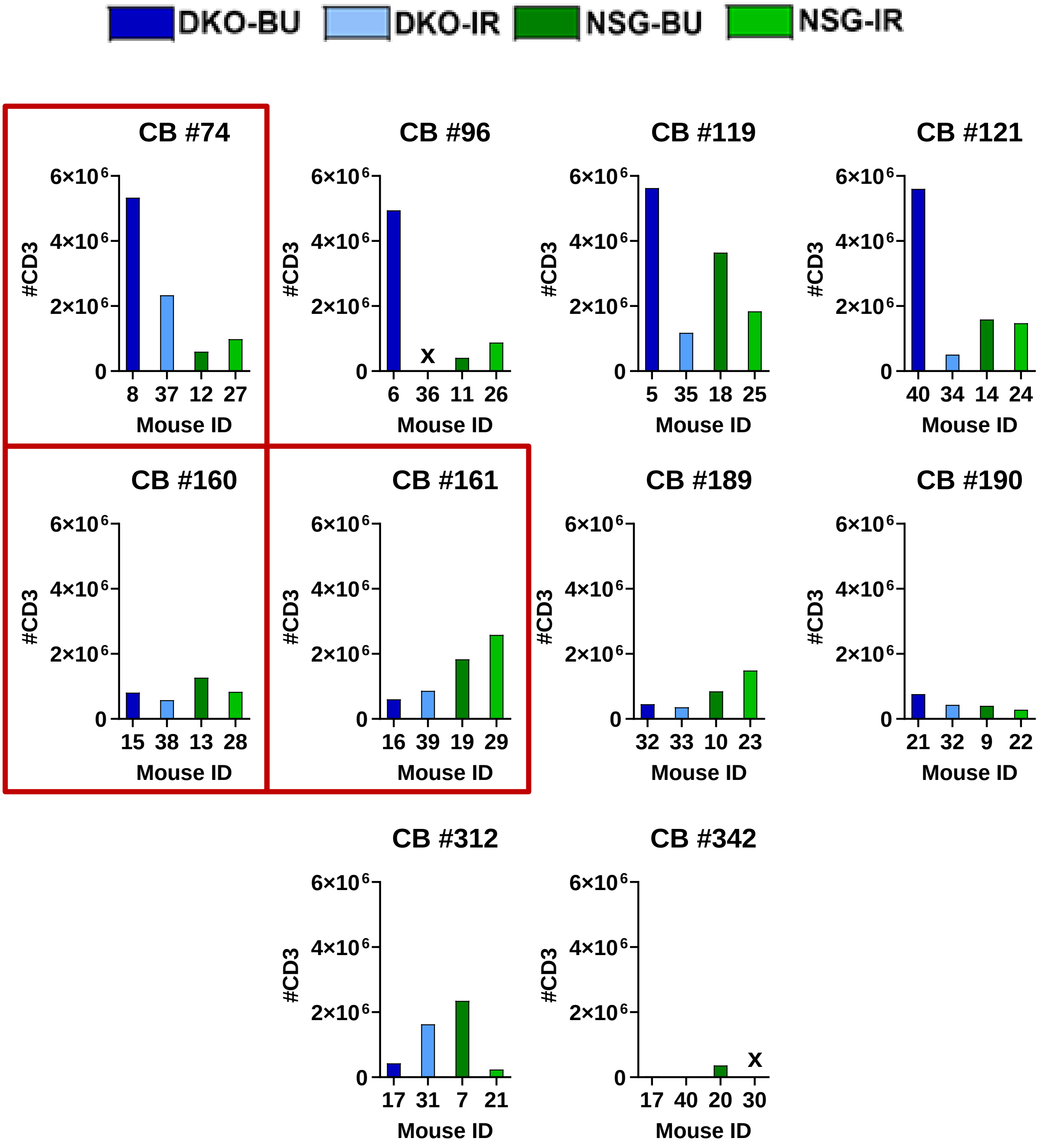

Figure S4. Marker gene expression used for annotation of splenic T-cell clusters.

A. Dot plot showing expression of selected marker genes across $CD4^{+}$ T cell clusters identified in Fig. 4, used for subset annotation. Genes are grouped into functional categories including naïve/memory, effector, and cytokine-associated markers.

B. Dot plot showing expression of selected marker genes across $CD8^{+}$ T cell clusters identified in Fig. 4, used for subset annotation. Genes are grouped into naïve/memory, effector, tissue-residency, cytokine, and exhaustion-associated markers.

In both panels, color indicates the average expression level of each gene within the cluster, and dot size represents the percentage of cells in the cluster expressing the gene.

**A**

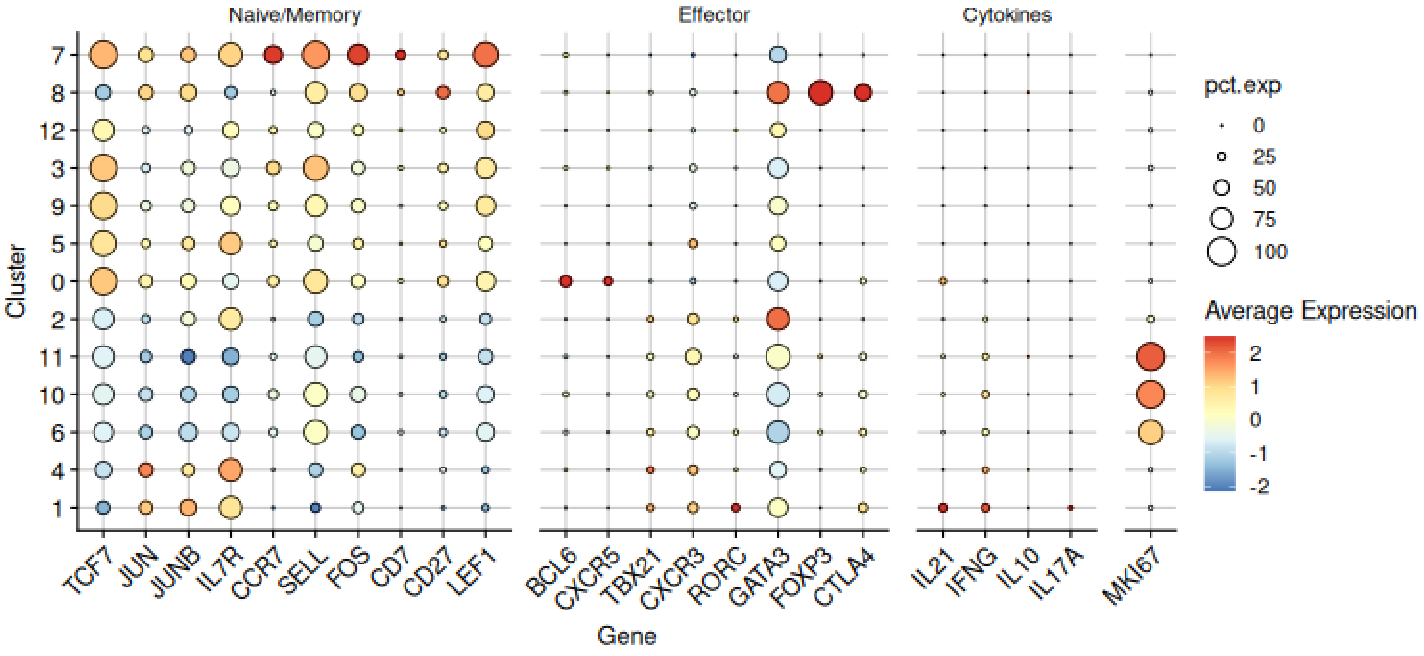


**B**

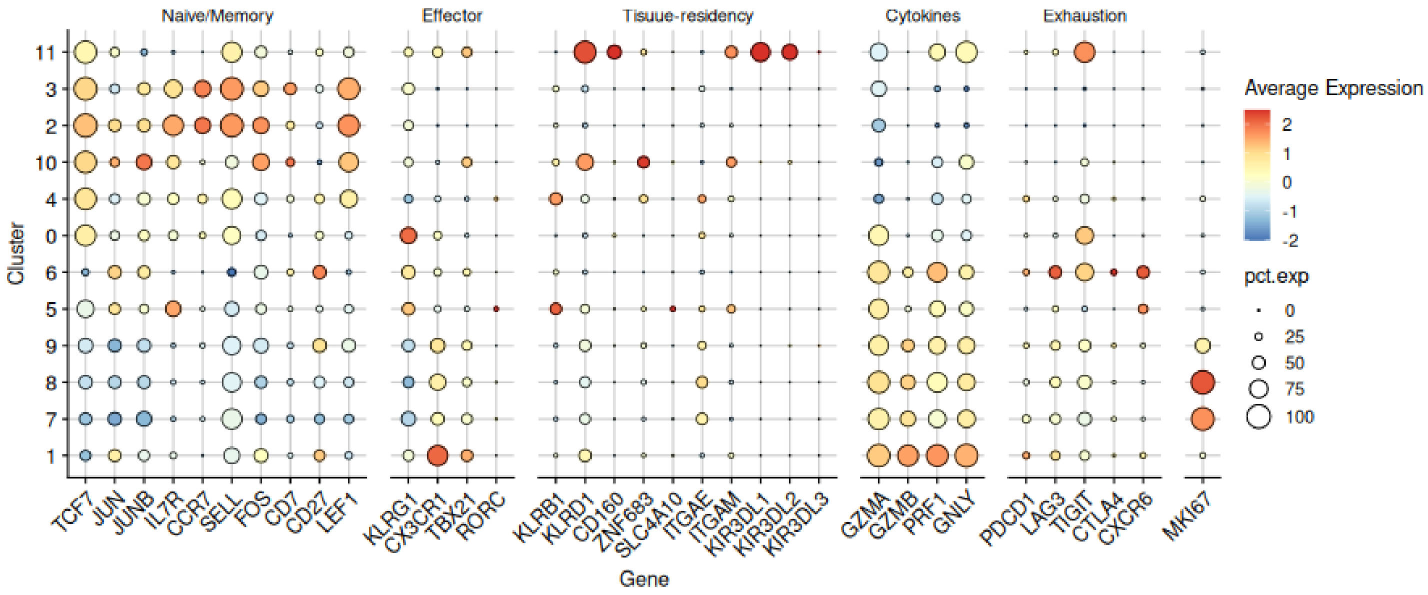

Figure S5. Distribution of T cell clone sizes across experimental cohorts and T cell subsets.
A, B. UMAP representation of $CD4^+$ T cells (A) and $CD8^+$ T cells (B) colored according to TCR clone size. Clones are categorized as small ($1\times10^{-4} < X \leq 0.001$), medium ($0.001 < X \leq 0.01$), or large ($0.01 < X \leq 0.1$) based on their relative frequency.
C. Bar plots showing the distribution of clone sizes across the four experimental cohorts (DKO BU, DKO IR, NSG BU, NSG IR) for $CD4^+$ T cells and $CD8^+$ T cells.
D. Bar plots showing the distribution of clone sizes across annotated T cell subsets for $CD4^+$ T cells and $CD8^+$ T cells. Clone size categories are indicated by color in all panels.

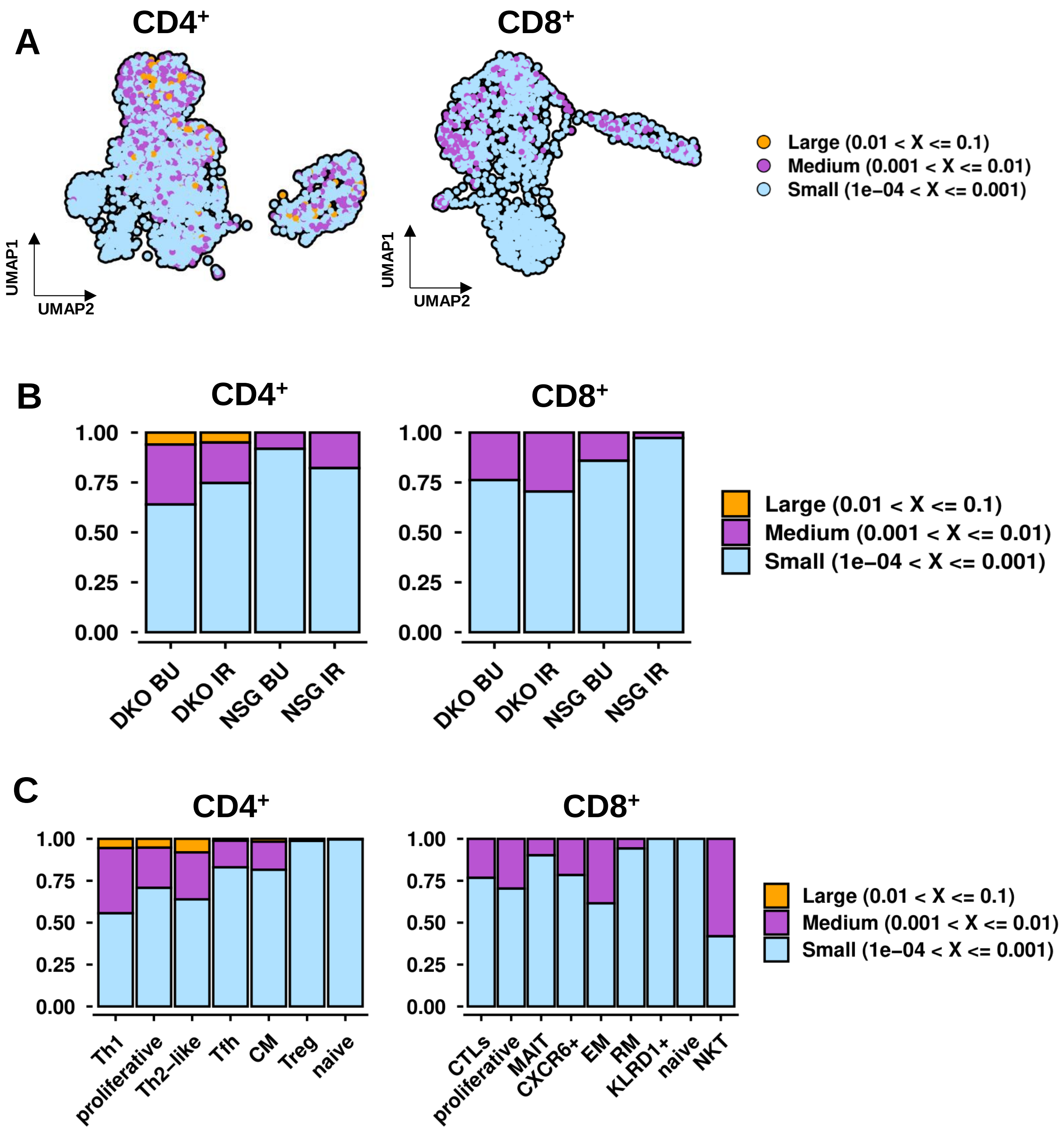

Figure S6: Data from TCR analysis of CD3$^+$ splenic cells.

A. Unique counts of TCR α, β, γ and δ within each cohort

B. Mean CDR3 legth TCR α, β, γ and δ within each cohort

C. Single cell mRNA seq analysis of CDR3 lenght within CD4$^+$ and CD8$^+$ splenic T cells.

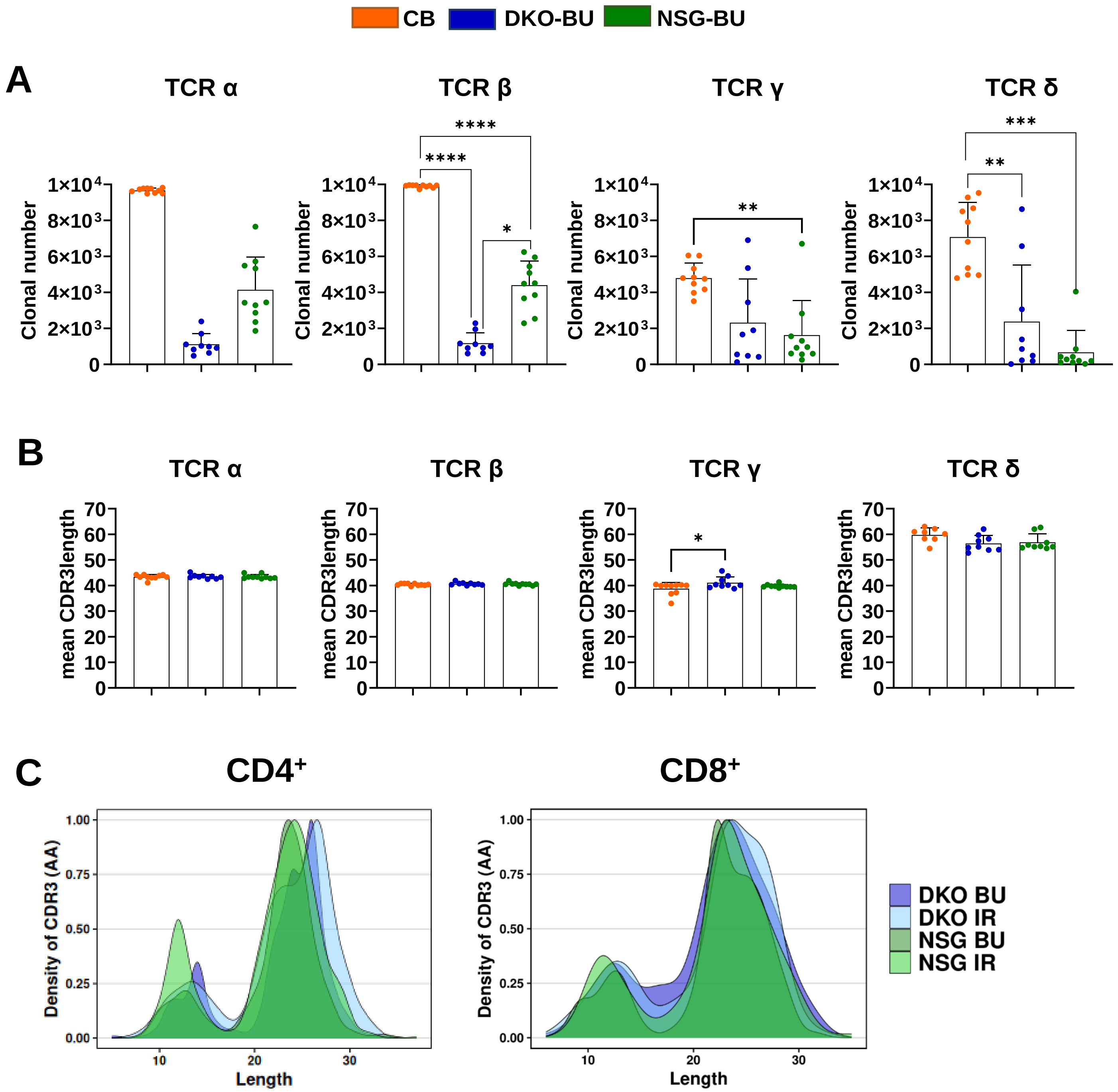

Figure S7: Heatmap visualization of T-cell receptor gene expression profiles in splenic CD3$^{+}$ lymphocytes derived from bulk TCR sequencing data:

A. TRA J genes

B. TRG V genes

**A**

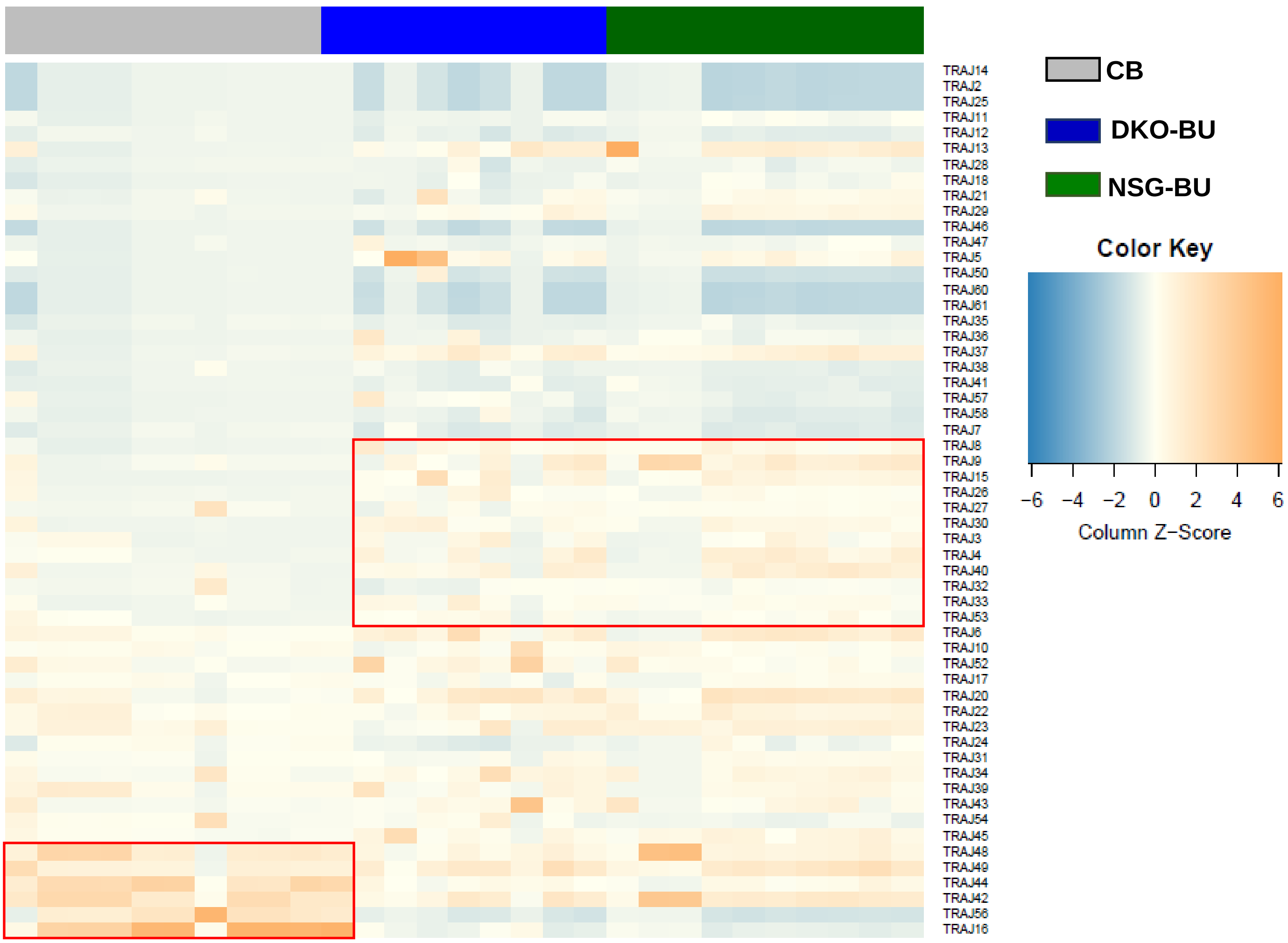


**B**

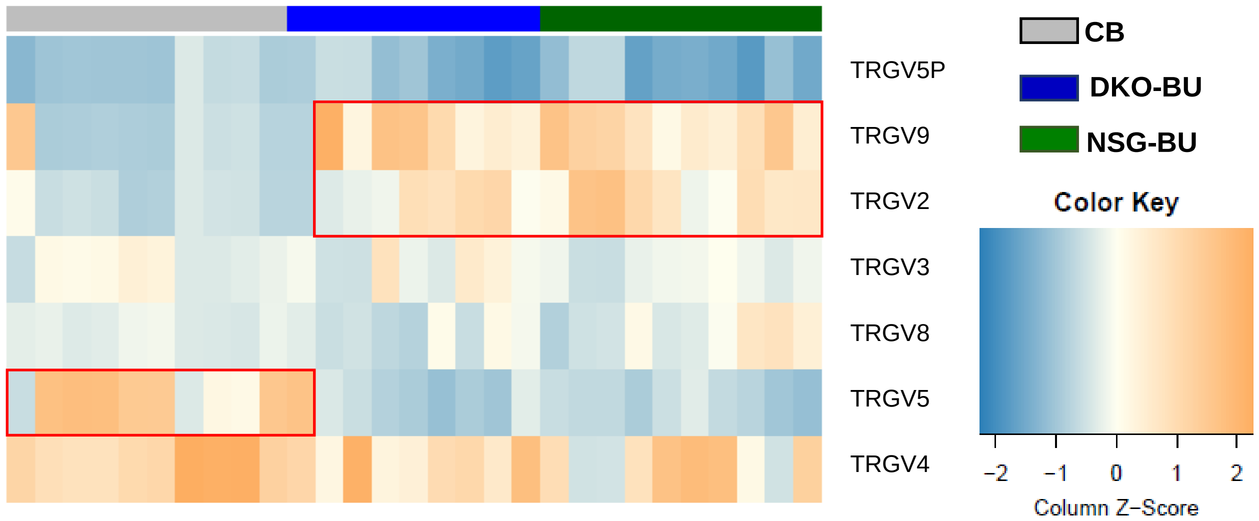

Figure S8: FACS Gating strategy for detection of human CD4$^+$ or CD8$^+$ T cells. Exemplary data for analyses of cells obtained from different sources.
A. Splenocytes of humanized mice
B. Human PBMCs
C. Human mononuclear cells from CD34$^-$ fraction cord blood
Cells were sequentially gated as follows: morphological selection on FSC vs SSC, doublet exclusion via FSC-H vs FSC-A, and viability discrimination. CD45 expression was used to distinguish human (huCD45$^+$) from mouse (mCD45$^+$) leukocytes. Within the huCD45$^+$ population, T cells (CD3$^+$) and B cells (CD19$^+$) were identified. CD3$^+$ T cells were further resolved into CD4$^+$ and CD8$^+$ subsets

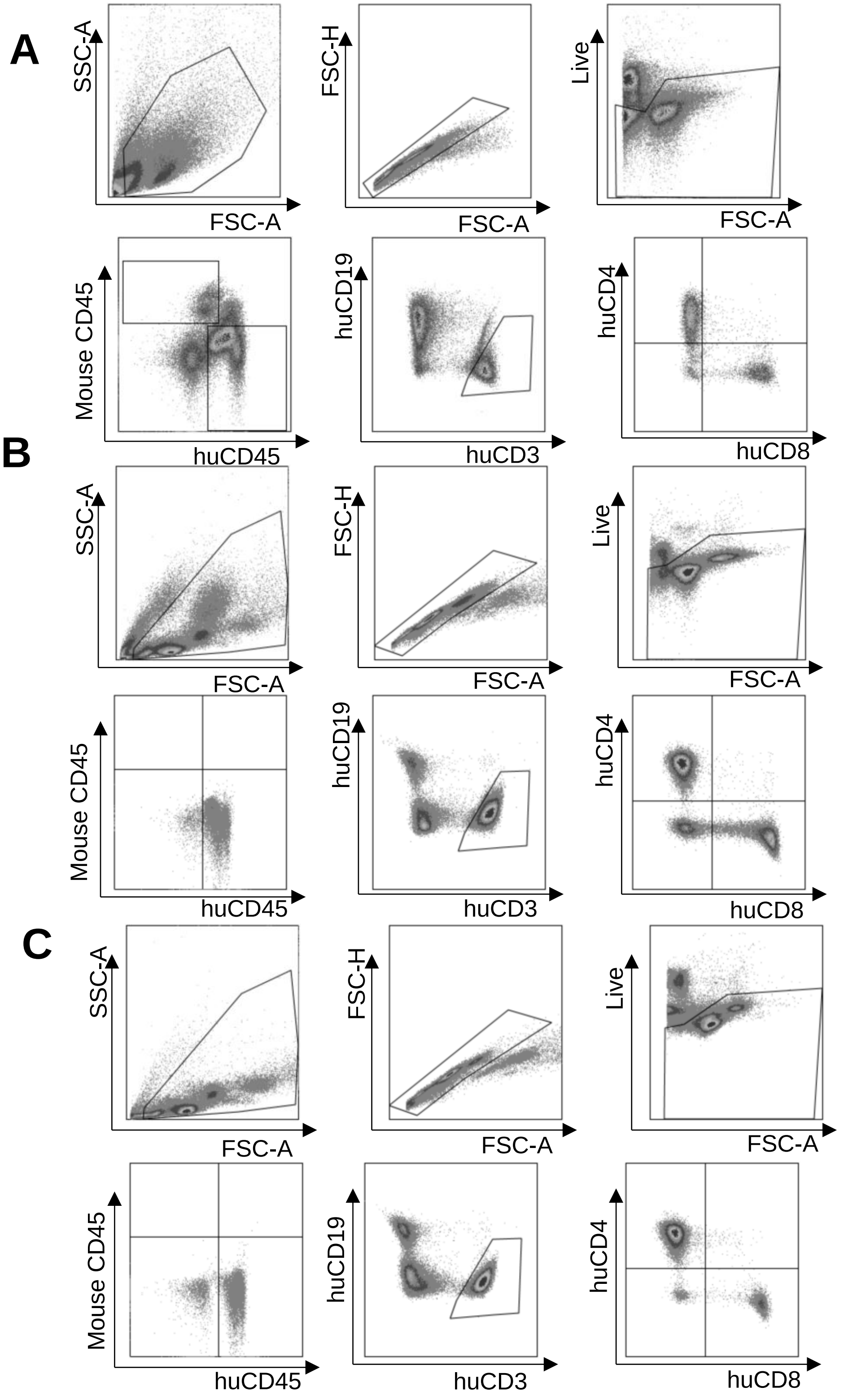

Figure S9: Gating strategy to quantify T cell immunophenotypes as naive (N), central memory (CM), effector memory (EM) or terminal effector (TE) via flow cytometry using cells obtained from different sources. CD4 cells shown in the left panels, CD8 cells shown in the right panels.
A. Splenocytes of humanized mice
B. Human PBMCs
C. Human mononuclear cells from $CD34^-$ fraction cord blood

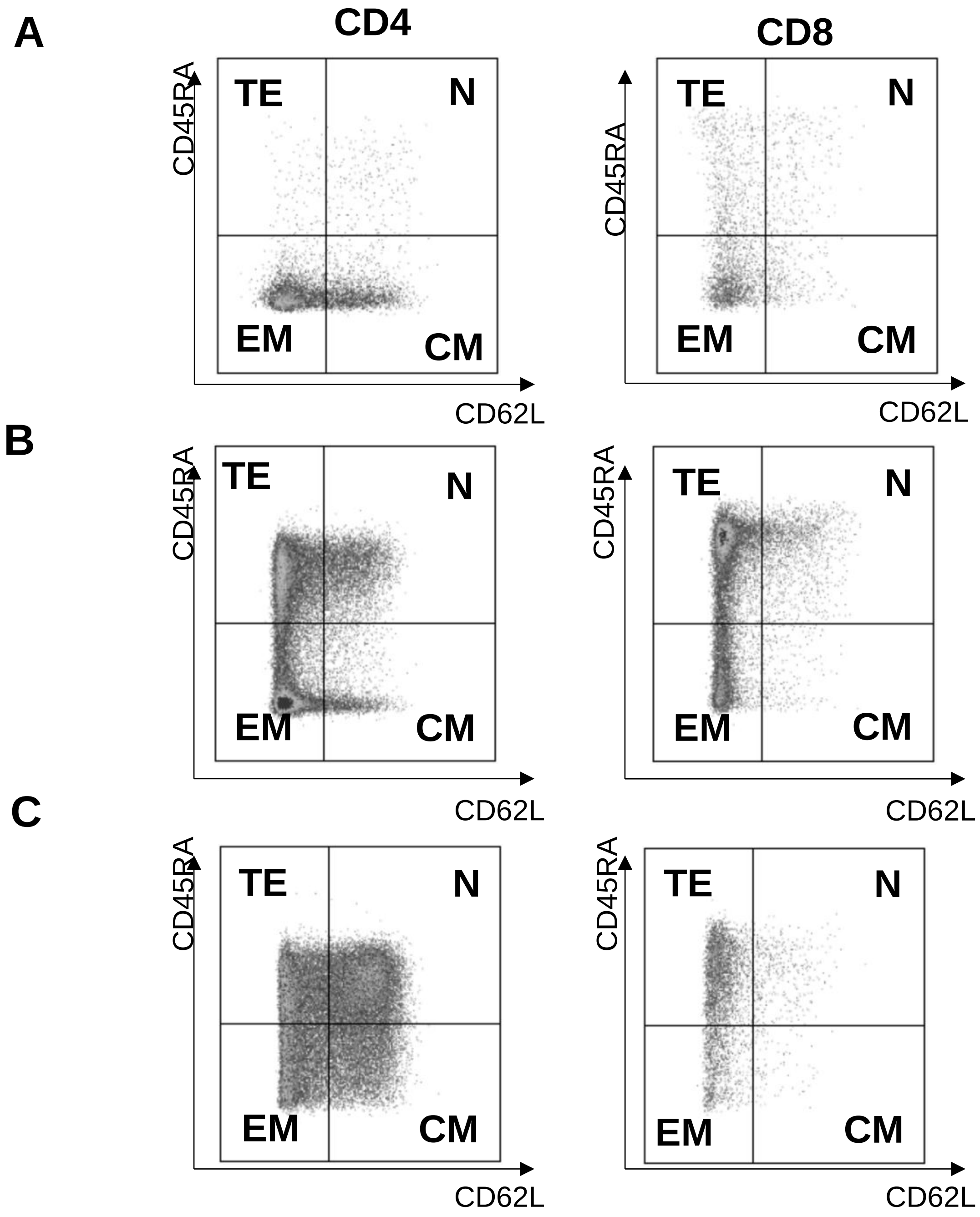
